# Leveraging three-dimensional chromatin architecture for effective reconstruction of enhancer-target gene regulatory network

**DOI:** 10.1101/2021.03.01.432687

**Authors:** Elisa Salviato, Vera Djordjilović, Judith M. Hariprakash, Ilario Tagliaferri, Koustav Pal, Francesco Ferrari

## Abstract

A growing amount of evidence in literature suggests that germline sequence variants and somatic mutations in non-coding distal regulatory elements may be crucial for defining disease risk and prognostic stratification of patients, in genetic disorders as well as in cancer. Their functional interpretation is challenging because genome-wide enhancer-target gene (ETG) pairing is an open problem in genomics. The solutions proposed so far do not account for the most updated knowledge on chromatin three-dimensional (3D) architecture, which is organized in a hierarchy of structural domains.

Here we introduce a paradigm shift based on the definition of multi-scale structural chromatin domains, integrated in a statistical framework to define ETG pairs. In this work *i*) we develop a computational and statistical framework to reconstruct a comprehensive ETG regulatory network leveraging functional genomics data; *ii*) we demonstrate that the incorporation of chromatin 3D architecture information improves ETG pairing accuracy; and *iii*) we use multiple experimental datasets to extensively benchmark our method against previous solutions for the genome-wide reconstruction of ETG pairs. This solution will facilitate the annotation and interpretation of sequence variants in distal non-coding regulatory elements. We expect this to be especially helpful in clinically oriented applications of whole genome sequencing in cancer and undiagnosed genetic diseases research.

## INTRODUCTION

Distal non-coding regulatory elements (enhancers) are crucial players in the control of gene expression. These are also the genomic features carrying the most marked epigenetic differences across cell types, thus constituting a fundamental component of the molecular and genetic mechanisms defining cell identity (1, 2). Enhancer activity status is itself regulated by epigenetics, chromatin accessibility and its three-dimensional (3D) architecture (3). In fact, the formation of chromatin loops allows distal regulatory regions to come in close physical proximity to their target gene promoters to regulate transcription (4). Their importance for human physiology is attested by their enrichment in polymorphisms associated to genetic diseases and cancer risk (5, 6). More mechanistic studies have shown the functional role of enhancer alteration in several pathologies, sometime collectively termed enhanceropathies (7, 8). Therefore, a genome-wide definition of the regulatory network constituted by enhancers and their target genes would be a valuable resource in biomedical research. For example, it would be instrumental for the annotation and interpretation of non-coding somatic mutations or germline sequence variants, to understand their effect on the broader gene regulatory network, in basic biology as well as in more translational studies.

Despite its importance, the reconstruction of a comprehensive network of enhancer-target gene (ETG) pairs remains elusive, especially because enhancers position with respect to the target genes is highly variable. Indeed, they can regulate one or more genes that appear distant in the linear sequence of the genome but may be in close physical proximity in the 3D chromatin organisation (9).

In this context, the development of molecular biology methods to study the 3D chromatin organization has been pivotal for achieving a better understanding of distal regulatory elements. In particular, the methods based on ligation by proximity, i.e. Chromosome Conformation Capture (3C) (10) and its high-throughput derivatives (11–13) (*e*.*g*. 4C, 5C and Hi-C), allow quantifying the frequency of physical interactions between distant chromatin regions (chromatin loops). Hi-C is the high-throughput genome-wide version of this technique, allowing researchers to map the contact frequency between virtually any pair of genomic loci (14).

In principle, Hi-C data could be used for the genome-wide identification of specific points of contact, such as ETG loops. However, Hi-C data is generally analysed by binning read counts at a resolution of few kilobases (kb), with the highest coverage datasets available to date reaching 1 kb (15–17). This resolution level is lower than what is needed to map ETG pairs when multiple enhancers are close to each other, or close to promoters. In all these cases, a distance smaller than 2 bins would not allow discriminating the interacting partners. Even the most recent literature, based on ENCODE3 data, reported that using Hi-C interaction calls to directly map ETG contacts is not a valuable strategy (18) to annotate distal regulatory elements, due to the resolution limit. Another challenge in this approach, is that different algorithms for calling point interactions in Hi-C data have generally discordant results and are influenced by the sequencing coverage (19).

Chromatin 3D architecture has not been optimally incorporated in the ETG network reconstruction algorithms proposed in literature so far. Some publications marginally used Hi-C data to call point interactions to be used as true positive contacts (20–23), despite the resolution and methodological shortcomings discussed above. Moreover, these approaches have been applied to a limited number of cell types, due to the reduced availability of Hi-C datasets. Alternatively, chromatin structure has been used only to restrict the initial search space of ETG pairs (24–27).

We hypothesised that the incorporation of experimental data on chromatin 3D architecture would enhance the accuracy of genome-wide ETG pairs definition. To this concern, here we introduce a paradigm shift based on the most up to date biological knowledge. Namely, there is a general consensus in the field about enhancer-target gene interactions occurring within the insulated boundaries of the so-called Topologically Associated Domains (TADs) (28, 29), which are relatively insulated domains enriched in local interactions. Moreover, several studies reported that TADs are largely conserved across different cell types (30–32). On the other hand, it is generally accepted that TADs can be defined at different levels of resolution, *i*.*e*. there is a hierarchy of TADs (17, 33, 34). More recent literature indicates that alternative TADs structures may indeed co-exist within a cell population, and the stochastic dynamics of active loop extrusion mechanisms could explain their formation and the patterns detected in Hi-C data (35–37). Therefore, we use multi-resolution TAD definitions as prior-knowledge to inform the selection of ETG pairs.

In this work 1) we develop a computational and statistical framework to reconstruct a comprehensive ETG regulatory network leveraging functional genomics data. Namely, we use a large panel of epigenomics datasets to define enhancer activity across multiple cell and tissue types, along with high resolution Hi-C data. 2) Then we demonstrate that the incorporation of chromatin 3D architecture information improves the accuracy in defining ETG pairs. In this context, we compute a score encoding the multiscale hierarchical structure of chromatin and use it as side information for controlling false discoveries and achieving high statistical power. 3) Finally, we extensively benchmark our method against previous solutions for the genome-wide reconstruction of ETG pairing. We show that our method is a valuable general-purpose solution, providing good ETG pairing performances for both long- and mid-range interactions.

## MATERIALS AND METHODS

### Definition of reference set of enhancer and gene promoter regions

We defined a reference set of enhancer regions using epigenomics datasets based on high-throughput sequencing across a compendium of cell and tissue types. In particular, we used ChIP-seq (Chromatin ImmunoPrecipitation followed by high-throughput sequencing) data for specific histone modifications, as detailed in the relevant results sections, as well as chromatin accessibility data based on DNase I hypersensitive sites (DHS), identified with DNase-seq. We downloaded histone H3 lysine 27 acetylation (H3K27ac) ChIP-seq and DHS narrow peaks (based on MACS v2.0.20 calls by the Roadmap Epigenomic consortium) called for 44 uniformly processed and consolidated cell and tissue types from Roadmap Epigenomic portal (https://egg2.wustl.edu/roadmap/data/byFileType/peaks/consolidated/narrowPeak/). H3K27ac is a post-translational histone modification associated with active enhancer and promoter regions, whereas DNase-seq allows assessing chromatin accessibility. We focused on the subset of cells and tissue types for which both H3K27ac ChIP-seq and DNase-seq were available (Table S1A). We further filtered the results for subsequent analyses considering only peaks with strong significant enrichment, *i*.*e*. −log_10_(adj.pvalue)*≥*5. Both the number (Figure S1A) and size (Figure S1B) of DNase-seq and H3K27ac ChIP-seq peaks vary across cell and tissue types. Namely, DNase-seq peaks were 123,400 on average, with average size 358bp. Conversely, H3K27ac peaks were fewer (53,721 on average) and larger (940 bp average size).

To obtain a comprehensive list of cis-regulatory elements we conducted a two-step procedure. Firstly, for each cell type, the intersection between H3K27ac and DHS peaks with overlapping regions (*≥*1 bp) were used to define cell-specific enhancers. Additional filters were applied ex-post, such as the removal of interval portions overlapping annotated exons (for both coding and non-coding genes) and the removal of intervals shorter than 10 bp or larger than 2.5 kb.

Secondly, cell-specific enhancers with overlapping intervals across different cell types were merged (union) together to define a consensus set of enhancer regions. This set was further annotated with respect to the transcription start site (TSS) as promoter-proximal (within 3.5 kb upstream and 1.5 kb downstream of TSS) or distal, and only the promoter-distal ones were retained as reference list of enhancer elements, hereinafter referred as *enhancer catalogue* (*n* = 347,777). This is meant to be a comprehensive reference set of regulatory regions, that can be active enhancers in at least one of the cell and tissue types considered. In the subsequent analyses to identify ETG pairs, we actually focused on the epigenetic status of the gene promoter regions, used as a proxy for the activity of the target genes. Thus, hereinafter we will refer to enhancer-promoter (EP) pairs when explicitly focusing on these genomic regions or epigenetic features, whereas we will refer to ETG pairs when focusing on the functional interaction to regulate the target gene. We defined reference promoters as 2 kb regions (1.5 kb upstream and 0.5 kb downstream) around the transcription start site (TSS) of annotated protein coding genes, based on RefSeq annotations in UCSC (refGene.txt.gz, May 2019, hg19 genome assembly). Non-canonical and Y chromosome were excluded. To reduce possible ambiguities, in case of multiple alternative transcripts for the same gene, only the most upstream TSS was maintained as reference for each gene. Moreover, promoters were merged in case of two close TSSes (±0.5 kb interval), *e*.*g*. divergent transcripts on opposite DNA strands, so as to obtain the final reference set of promoter regions (*m* = 18,027).

To compare our enhancer catalogue with alternative functional genomic definitions we employed: i) the atlas of active enhancers provided by FANTOM5 project (https://fantom.gsc.riken.jp/5/data/) (38), based on 808 human Cap Analysis of Gene Expression (CAGE) experiments (39); and ii) the collection of *in vivo* validated enhancers coming from the VISTA Enhancer database (40), based on transgenic mice reporter assays in 23 tissues of mouse embryos (41). We downloaded enhancer coordinates from FANTOM5 repository (https://fantom.gsc.riken.jp/5/datafiles/latest/extra/Enhancers/human_permissive_enhancers_phase_1_and_2.bed.gz), and we retrieved positive (*i*.*e*., elements that show consistent reporter gene expression among at least three embryos) “Human only” enhancers from VISTA Enhancer Browser (date version: 12 February 2020). In line with the procedure used to define our enhancer catalogue, interval portions overlapping annotated exons and promoter proximal elements were removed to obtain the final set of enhancers from each of these alternative sources. These filtered FANTOM and VISTA enhancer sets were used in the subsequent analyses and are composed of 58,200 and 894 enhancers, respectively.

### Hi-C dataset processing

We leveraged chromatin 3D architecture data from genome-wide chromosome conformation capture experiments based on high-throughput sequencing (Hi-C). Namely, we processed eleven Hi-C datasets (Table S1B) covering different cell lines and primary tissues from a compendium of public datasets (17, 42–46).

For each Hi-C dataset we retrieved the raw FASTQ files from the NIH SRA database (https://www.ncbi.nlm.nih.gov/sra). The sequencing reads were aligned with the iterative mapping procedure (single-end mode) as implemented in hiclib (https://github.com/mirnylab/hiclib-legacy)(version from gitHub commit d38f198, date: September 28, 2017) based on botwie2 (version 2.3.4.3) aligner (47) (https://github.com/BenLangmead/bowtie2). Briefly, in this iterative alignment procedure reads were truncated at 30 bp and aligned to the reference genome (hg19). Reads that were not uniquely aligned were elongated (5 bp) and the alignment procedure repeated, with additional iterations until full read length or successful alignment is achieved. For each FASTQ file the information on uniquely mapped reads were stored in a HDF5 (Hierarchical Data Format) file. Biological or technical replicates belonging to the same dataset were merged in a single HDF5 file (hdf5 library, version 2.9.0). We filtered read pairs with a sum of distances from the downstream restriction site not compatible with the expected fragment size: *i*.*e*., events originating from non-canonical enzyme activity or non-enzymatic physical breakage. The distance cut-off was estimated for each dataset based on the frequency distribution of distances and the expected fragment length. Duplicated read pairs, as well as read pairs derived from unligated or circularized fragments, were also removed.

Finally, the genome was binned at 10 kb bin size, and the raw read counts were summarized in a Hi-C contact matrix for each chromosome, accounting for intra-chromosomal interactions. To allow comparability among all tissues and cell types and correct for technical biases, chromosome-wise iterative correction (ICE) with default parameters (48, 49) was applied (using cooler version 0.8.5, https://github.com/open2c/cooler). This procedure returned a balanced matrix of relative contact probabilities, in which each row (excluding the elements in the first two removed diagonals) summed up to one. The output files (cool format) were converted to txt files and compressed.

### Hierarchical Contact Likelihood score

To account for the three-dimensional spatial proximity of regulatory elements, we devised a score proportional to the likelihood of enhancer-promoter (EP) pairs co-localization, named Hierarchical Contact Likelihood (HCL) score. HCL accounts for the TADs hierarchical structure across multiple tissue and cell types. For HCL definition we relied on the Local Score Differentiator (LSD) TAD borders calling procedure (50), as implemented in the HiCBricks (version 1.8.0) Bioconductor package (51). We defined TADs as regions between two consecutive domain boundaries. LSD is based on the directionality index (DI) score originally proposed by Dixon et al. (30). Among the user defined parameters in this algorithm, the DI-window (*i*.*e*., the number of up-stream and down-stream bins over which the DI score is computed) influences the scale of the TAD domains that are identified: the larger the DI-window, the larger the average resulting TAD size.

The HCL score is thus defined by considering a collection 𝒟 of Hi-C contact matrices (all binned at the same bin size) and an ensemble of TADs boundaries for each D ∈ 𝒟, denoted by TAD_D_(w), where w ∈ W is the DI-window. We considered *W* = [5, 10, 20, 50] for the DI-window size. Thus, each TAD_D_(W) represents the multi-resolution TADs structure for a specific Hi-C contact matrix in a specific cell and tissue type.

Given a list of n enhancers and m promoters, we can define a binary matrix M_D_(w), in which an element (i, j) is set to 1 if the i-th enhancer and the j-th promoter are within the same TAD belonging to the ensemble TAD_D_(w). The matrix M_D_(w) is thus an n × m co-occurrence matrix. To estimate the overall spatial relationships between enhancers and promoters over the hierarchical structure of each Hi-C contact matrix D, we propose the aggregate score:

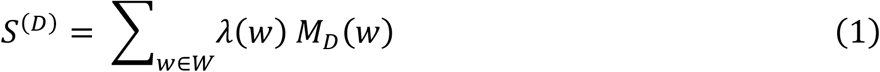

where 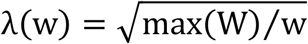 is a scaling factor that gives higher weight to smaller TAD hierarchies according to the set W. Namely, we are setting the highest level to have a weight equal to one. To extend the score to the entire collection 𝒟 of Hi-C matrices, we define the HCL score as:

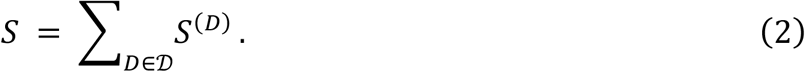

Each element of the matrix *s*_ij_ is meant to capture the broader spatial co-localization pattern of the i-th enhancer and the j-th promoter across both different layers of TADs hierarchy and tissue or cell types. By definition, for the set *W* = [5, 10, 20, 50] and using 11 Hi-C datasets, the lower and upper limits of the score are min(*S*) = 0 and max(*S*) = 87.8, respectively, where the maximum for each Hi-C contact matrix is equal to max (*S*^*(D)*^) = 7.98.

As the HCL score can be calculated whenever a hierarchy of TADs is provided, for comparison purposes we considered TopDom (52), a public available tool meant to identify TADs at sub-mega base resolution. TopDom identifies TAD boundaries looking at significant local minima of the bin signal function, which is computed with a procedure similar to the previously proposed insulation score (53). Namely, the bin signal function is the average contact signal in the neighbourhood of each bin along the diagonal, considering a diamond-shape area of with 2*ŵ* where *ŵ* is a tuneable parameter that defines the window size. We used TopDom R package (https://github.com/HenrikBengtsson/TopDom, version 0.8.1) to call TADs, defined as regions between two boundaries flagged as significant minima (local.ext=−1). For the *ŵ* parameter we used the same set of values adopted for LSD DI-window sizes, i.e. *W* = *Ŵ* = [5,10,20,50].

### Enhancer-promoter pairs synchronization analysis with Canonical Correlation

We adopted the Canonical-Correlation Analysis (CCA) (54) to quantify the strength of coordinated activity in each EP pair. We considered enhancer and promoter regions separately, and quantified their respective activity status using two sets of epigenetic marks: we used the enrichment of DNase-seq and H3K27ac ChIP-seq (*p* = 2) for enhancers and DNase-seq, H3K27ac and H3K4me3 (*q* = 3) for the promoters.

Namely, we downloaded H3K27ac, H3K4me3, and DNase-seq consolidated fold-change enrichment signal tracks (bigwig format) from the Roadmap Epigenomic consortium web portal (https://egg2.wustl.edu/roadmap/data/byFileType/signal/consolidated/macs2signal/foldChange/) for all the cell and tissue types for which all the three epigenetics marks were available (*k* = 44) (Table S1A). For each enhancer and promoter region, we computed the maximum signal from the proper bigwig genomic tracks, using rtracklayer R package (version 1.44.4) (55).

We then used CCA to investigate the inter-set correlation patterns. More formally, let *X*^(*i*)^ denote a *p*-dimensional random vector of quantitative features describing the activity of the *i*-th enhancer. Let *Y*^*(*j*)*^ denote a *q*-dimensional random vector of quantitative features describing the activity of the *j*-th promoter. Our data consist of *k* independent observations of *X*^(*i*)^ and *Y*^*(*j*)*^ across *k* cell and tissue types. We are interested in testing the null hypothesis of independence between *X*^(*i*)^ and *Y*^*(j)*^, *i*.*e*., the lack of synchronized activity:

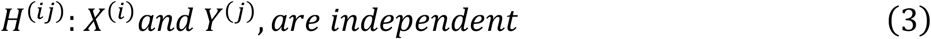

against a general alternative. Assuming normality of *X*^(*i*)^ and *Y*^*(j)*^, the null hypothesis of interest can be equivalently expressed as:

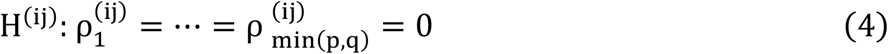

where 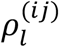 is the *l*-th canonical correlation coefficient associated to *X*^(*i*)^ and *S*^(*j*)^. Briefly, the canonical correlation coefficients measure the correlation over subsequent linear transformations of the original *p* and *q* variables, that allow maximizing the relationship between the two sets, while ensuring independence within each set. The maximum number of linear transformations is min(p, q), *i*.*e*., two in our case. The key advantage of CCA is to reduce the dimensionality and the inter-confounding factors of each set, while extracting the major correlation patterns.

Following the CCA, we calculate the p-value for the null hypothesis (4) by testing the sequential hypotheses that the first canonical correlation coefficient, and all the following ones, are zero using the Wilk’s lambda statistics (54):

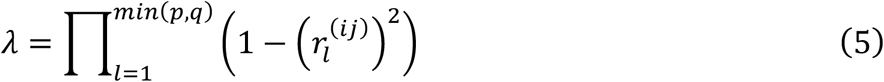

where 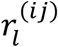 is the estimated *l*-th canonical correlation coefficient. To improve the accuracy for small sample sizes, we adopted the Rao’s F-approximation (56). Namely, *λ* was transformed to an j-statistic using Rao’s approximation as implemented in the candisc R package (version 0.8-3).

The procedure returned a single p-value *p*_(*ij*)_ for the overall dependence of the j-th promoter on the j-th enhancer. *p*_(*ij*)_ quantifies the amount of evidence provided by the data for the presence of the synchronized activity between a specific EP pair.

### 3D architecture integration in the enhancer-promoter pairs FDR control

The reconstruction of the EP pairs based on CCA, as described above, is based on testing millions of hypotheses (*i*.*e*., one for each EP pair), thus requiring some control over the number of false discoveries. In large scale testing problems of this kind, the typical goal is the control of the False Discovery Rate (FDR), defined as the expected fraction of false discoveries. The Benjamini-Hochberg (BH) (57) correction is a frequently used method for controlling FDR in genomics data analyses. In this work, however, we aim to increase statistical power over the standard BH procedure by considering relevant contextual information. An example is provided by the 3D co-localization information encoded in the HCL score. To include this information in the testing problem, we relied on the Adaptive *p*-value thresholding procedure (AdaPT), recently proposed by Li and colleagues (58) and implemented in the adaptMT R package (version 1.0.0). AdaPT estimates Bayes-optimal *p*-value rejection threshold based on user-defined side information, and controls FDR in finite samples. Five logistic-Gamma generalized linear models with natural cubic splines as candidate models were explored to identify the best threshold estimate, as proposed by default parameters of AdaPT implementation.

Formally, we considered our collection of hypotheses {*H*^(*ij*)^ *i* = 1,… *n, j* = 1,… *m*}, as defined in (4) for which we computed i) a *p*-value *p*^(*ij*)^ ∈ [0,1] quantifying the strength of evidence for the presence of a synchronized activity between the *R*-th enhancer and the *T*-th promoter and ii) a score *s*_*ij*,_ ∈ ℝ capturing the likelihood of their 3D proximity. Then, we used AdaPT to determine a rejection threshold in function of the HCL score, *i*.*e*., *f*(s), such that the estimate of the False Discovery Proportion (FDP) is bounded by a prespecified level *α* ∈

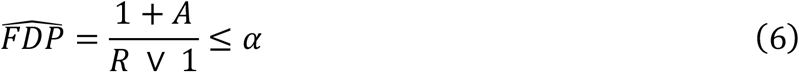

Where *R* = |{*p*^(*ij*)^: *p*^(*ij*)^ ≤ *f*(*s*_*ij*_) | and *A* = |{*p*^(*ij*)^: *p*^(*ij*)^ > 1 − *f*|(*s*_*ij*_)} | are the number of rejected and not rejected null hypotheses, respectively. By Theorem 1 of (58), the associated FDR is also bounded by α. AdaPT has been shown (58, 59) to significantly increase statistical power in situations in which the considered side information provides a useful basis for prioritizing most promising hypotheses. Nevertheless, statistical guarantees regarding FDR control are preserved also when the side information is inaccurate or not relevant for the problem at hand: in this case, the weight given to the side information will be low and AdaPT will converge to the standard BH method.

### Reference benchmarking datasets

#### Expression quantitative trait loci databases

Expression quantitative trait loci (eQTLs) are Single Nucleotide Polymorphisms (SNPs) associated to an alteration in the expression of a specific gene. We considered multiple eQTL datasets as reference for benchmarking the pairing of distal regulatory elements to their target gene. In particular, we considered eQTL data from i) the Genotype-Tissue Expression (GTEx) project (60), with eQTLs inferred from a panel of 15,201 samples in 48 tissue types; and ii) the pan-cancer eQTL (PanCanQTL) analysis (61), with eQTLs inferred from a panel of 9,196 tumour samples in 33 cancer types from The Cancer Genome Atlas (TGCA).

Cis-eQTL files from the v8 GTEx data release for 48 tissue types (Table S1C) were downloaded from GTEx portal (https://www.gtexportal.org/home/datasets). All *.sign_variant_gene_pairs.txt.gz files were converted to GenomicRanges (1.36.1, Bioconductor package) objects and merged maintaining only one eQTL in case of redundancy. The *.egenes.txt.gz files were used to convert Ensemble gene IDs to Gene Symbols. Genomic coordinates were converted from hg38 to hg19 genome build using liftOver tool (rtracklayer R package version 1.44.4 (55)). Cis-eQTL files from PanCanQTL (Table S1D) were downloaded from URL http://gong_lab.hzau.edu.cn/PancanQTL/cis (filenames with suffix *_tumor.cis_eQTL.xls). All Cis-eQTL files were converted in GenomicRanges objects and merged maintaining only one eQTL in case of redundancy.

An EP pair was considered supported by an eQTL (*i*.*e*., validated) if the corresponding SNP was located within an enhancer genomic region and associated with the expression of the cognate promoter. If multiple SNPs were within the same enhancer genomic region, they were considered only once. If the same eQTL was predicted in multiple tissue types to regulate a specific target gene, it was also considered only once.

#### Capture Hi-C datasets

We also considered nine capture Hi-C (cHi-C) experiments (Table S1E) coming from seven different studies (62–68), specifically designed to identify DNA-DNA interaction between promoters and distal chromatin regions. All the downloaded interaction lists (washU format) were already pre-processed in the original articles, and CHICAGO (Capture Hi-C Analysis of Genomic Organization) (64) algorithm was used to select significant interactions (− log_10_(adj.pvalue) *≥* 5). Genomic coordinates were converted from hg38 to hg19 genome build using liftOver tool (rtracklayer R package version 1.44.4 (55)), when needed.

An EP pair was considered supported by a cHi-C interaction if the promoter region overlaps (*≥*1 bp) with the “bait fragment” and the enhancer with the “other end fragment”, or vice versa. Ambiguous EP pairs due to the cHi-C resolution (*i*.*e*., pairs supported by the same cHi-C interaction) were not discarded.

#### BENGI benchmark

As an additional reference dataset, we considered the Benchmark of candidate Enhancer-Gene Interactions (BENGI) dataset (69). We downloaded *All-Pairs*.*Natural-Ratio* files from BENGI GitHub repository (https://github.com/weng-lab/BENGI/). This included a total of 21 lists of curated interactions (Table S1F) supported by ChIA-PET, Hi-C, eQTL and CRISPR genome editing experiments and covering seven cell lines and six tissue types. For each file, only the enhancer-like signatures (*i*.*e*., the ones marked as high DNase and H3K27ac signal) were considered. In line with the previous section (“Definition of enhancer and gene promoter regions”), but using BENGI gene definitions (GENCODEv19-TSSs.bed.gz annotation file), we defined promoter intervals as 2 kb windows (1.5 kb upstream and 0.5 kb downstream) around the transcription start site (TSS). Only the most upstream TSS for each gene was preserved. Enhancer intervals were annotated using the hg19-cCREs.bed.gz file. All 21 lists of enhancers and promoters were pooled to perform the EP pairing analysis based on our framework and then split for the assessment. An EP pair was deemed as true positive if supported by a specific BENGI curated interaction (*i*.*e*., the internal flag was equal to 1).

### ETG pairs by other tools

To benchmark our ETG pairing framework against other algorithms, we considered state-of-the-art methods among the 36 listed in a recent review (70). To overcome limitations related to the lack of user-friendly software, we considered only algorithms with publicly available ETG pairs lists, called as described in the original publications. Namely, the selection resulted in seven tools (Table S1G): FOCS (FDR-corrected OLS with Cross-validation and Shrinkage) (71), PreSTIGE (Predicting Specific Tissue Interactions of Genes and Enhancers) (72), RIPPLE (Regulatory Interaction Prediction for Promoters and Long-range Enhancers) (73), PETmodule (Predicting Enhancer Target by modules) (22), TargetFinder (23), JEME (Joint Effect of Multiple Enhancers) (21) and DeepTACT (Deep neural networks for chromatin conTACTs prediction) (74).

For each algorithm, we downloaded the lists of ETG pairs for all the available cell and tissue types and we processed them to obtain a uniform format of annotations. Namely, for each ETG pair we stored: i) enhancer genomic region coordinates (chr:start-end); ii) promoter region or TSS genomic coordinates (chr:start-end), depending on the information reported by the authors; iii) gene symbol; iv) prediction flag (*i*.*e*., 1: predicted, 0: not predicted), as sometimes the original authors reported only the predicted pairs and sometimes also the entire set of initial candidates; v) distance between enhancer mid-point and promoter mid-point (or TSS); and other optional information returned by the specific algorithm (*e*.*g*., score, etc). Genomic coordinates were converted from hg38 to hg19 genome build using liftOver tool (rtracklayer R package version 1.44.4 (55)), when needed. Gene symbol and TSS coordinates were retrieved from BioMart database, through R Bioconductor interface (version 2.40.5, host=“grch37.ensembl.org”, path=“biomart/martservice/”, database=“hsapiens_gene_ensembl”). ETG pairs associated to Ensemble gene IDs without any match with gene symbols were discarded. To make enhancers of the other tools comparable with our enhancer reference catalogue, we applied the filters described in section “Definition of reference set of enhancer and gene promoter regions” with minor modifications. Namely, we removed interval portions overlapping annotated exons (for both coding and non-coding genes, RefSeq annotations in UCSC) and promoter proximal elements. If one or more exons were completely located within an enhancer interval, the enhancer was split and the pair duplicated in concordance with the number of resulting enhancers. A promoter proximal element was defined as a pair with distance between enhancer mid-point and promoter mid-point (or TSS) smaller than 3 kb.

To evaluate the performances of our method in identifying cell type-specific ETG pairs, we performed a direct comparison with JEME, using the initial set of enhancers, gene and candidate ETG pairs, inferred from 127 cell and tissue types, collected by the Roadmap Epigenomic consortium (Supplementary Table S1F). In line with the previous section (“Definition of reference set of enhancer and gene promoter regions”), but using JEME gene annotations, we defined promoter intervals as 2 kb windows (1.5 kb upstream and 0.5 kb downstream) around the transcription start site (TSS). All lists of enhancers and promoters were initially pooled together to perform the EP pairing analysis based on our framework, and then split for the assessment. The HCL scores were calculated based on EP pairs colocalization and no cut-off was applied (i.e. we did not filter pairs with HCL >1). Candidate pairs were sorted based on the “confidence score” (descending order) and p-values (ascending order), for JEME and our framework, respectively.

To investigate the expression of the predicted target genes we downloaded all the available matched consolidated RNA-seq profiles (57 out of 127) from Roadmap Epigenomics portal (https://egg2.wustl.edu/roadmap/data/byDataType/rna/expression/57epigenomes.RPKM.pc.gz).

### Assessment of predicted ETG pairs and other indices

For each ETG calling algorithm (Table S1F) we define as *Z* the number of candidate pairs, *i*.*e*., the initial number of input EP pairs for which a score or *p*-value was calculated by the original authors; *V* the number of true pairs, *i*.*e*., all the EP pairs contained in at least one of the reference benchmarking datasets; *z* the number of predicted pairs, i.e., the EP pairs that satisfied the selection criteria as applied by the original authors (for example *p*-value ≤ α); *v* the number of true predicted pairs, i.e., the predicted EP pairs contained in at least one of the reference benchmarking datasets.

We used four different indices for performance assessment: *Precision* (P), the percentage of true predicted pairs over the total number of predicted pairs (*v*/*z*); *Recall* (R), the percentage of true predicted pairs over the total number of true pairs (*v*/*z*); *F1 score*, the harmonic mean of precision and recall 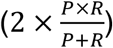 and *Relative Improvement* (RI), the improvement respect to random choice 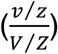 Precision-recall curves were computed by sorting EP pairs based on distance, HCL score, canonical correlation or HCL-based AdaPT corrected p-value, and calculating precision and recall for all the possible cut-offs (*Z*) of the candidate pairs list.

The Jaccard Index (JI) between two sets of genomic regions was calculated as i) the number of elements that overlap (bp>0) over the total number of elements in the two sets (*i*.*e*., JI on overlap); or ii) the total length of intersections divided by the total length of the union of the two sets (*i*.*e*., JI on coverage).

## RESULTS

### Methodological framework overview

Here we present a new approach for the definition of enhancer-target gene (ETG) pairs leveraging the most updated biological knowledge on chromatin 3D architecture and integrating heterogeneous functional genomics data into a rigorous statistical framework. Our method is introducing a general framework leveraging a paradigm shift in the integration of chromatin 3D architecture. Its three key features are:

#### 1) Statistical framework for quantifying enhancer-promoter pairs synchronization

The method is flexible in terms of input, as it starts from user-defined sets of i) enhancer and promoter regions and ii) functional genomics data to quantify their activity (Figure 1A). This flexibility is ensured by the use of Canonical-Correlation Analysis (CCA) to quantify the synchronization of enhancer-promoter (EP) pairs activity across cell types. Moreover, it is designed to leverage multiple types of functional genomics data, also accounting for the correlation within sets of features.

**Figure 1.**
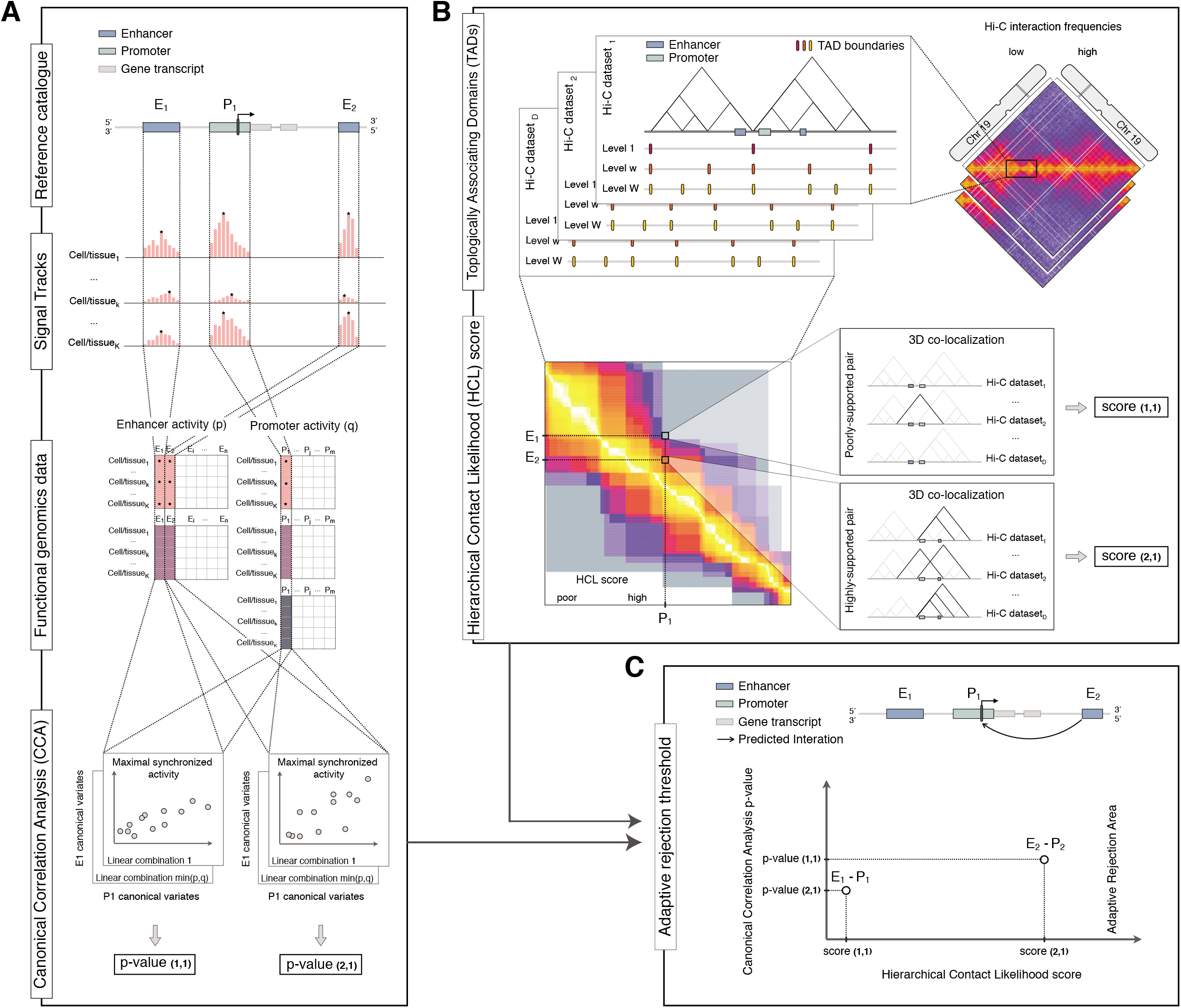
Methodological framework overview. The schematic illustration of the workflow of our methodological framework. **A**. Starting from a reference catalogue of enhancer and promoter regions, it is possible to quantify their respective activity status using two sets of *p* and *q* functional genomic data (e.g. ChIP-seq data for chromatin marks), respectively. Then, the Canonical Correlation Analysis (CCA) is used to investigate the synchronised activity of each enhancer-promoter (EP) pair across *k* cell and tissue types: the two original sets of chromatin marks are transformed through linear combinations that allow maximizing the relationship between the two sets, and the respective canonical correlation is tested. The procedure returns a *p*-value for each specific EP pair. **B**. For each Hi-C dataset in the selected collection, the boundaries of Topologically Associating Domains (TADs) are identified across multiple levels of resolution. The resulting ensemble of boundaries represents the hierarchical structure of TADs for a specific cell or tissue types. Considering the occurrence of each EP pair within these ensembles called from Hi-C datasets, we can describe their broader spatial co-localization pattern through the Hierarchical Contact Likelihood (HCL) score. A high score is associated to pairs supported by several combinations of Hi-C datasets and hierarchical levels (e.g., *E*_2_ − *P*_1_ pair). Conversely, a weak score is associated to pairs supported only in few combinations (e.g., *E*_1_ − *P*_1_ pair). **C**. The 3D co-localization information encoded in the HCL score is used to estimate an adaptive rejection threshold to control for FDR in the multiple testing hypothesis of EP pairs synchronisation. On similar equal nominal *p*-value (y-axis) a less conservative significance criterion is used for the EP pair showing higher HCL score (x-axis). Namely, even if one enhancer (*E*_1_) will exhibit a stronger synchronization with a specific promoter (*P*_1_), being at greater 3D distance will be less likely to regulate it than the closest one (*E*_2_).

#### 2) Hierarchical Contact Likelihood (HCL) Score

It takes into account the chromatin architecture as experimentally measured by Hi-C, to compute the HCL score accounting for the likelihood of ETG pairs 3D co-localization (Figure 1B). Differently from previous methods, we leverage up-to-date biological knowledge on TADs role in physically confining ETG pairs interactions, in addition to TADs multi-scale hierarchical organization and their conservation across cell types.

#### 3) Chromatin 3D architecture and functional genomics data integration

The information on chromatin 3D architecture is used to increase the statistical power to detect ETG pairs synchronization, while controlling false discoveries (Figure 1C). Previous literature primarily used chromatin conformation capture data to define point interactions and train machine learning classifiers, or to validate ETG pairs (70, 75). This is the first time that chromatin 3D architecture is directly integrated as side information in the statistical model for defining ETG pairs.

### Definition of the reference enhancer catalogue

The first challenge in the definition of ETG pairs is the lack of a universal reference list of enhancer regions, as they do not have a univocal nucleotide sequence. A comprehensive definition of enhancers based on functional genomics data in principle would require analysing virtually every cell and tissue type. This is practically impossible, despite ambitious large-scale collaborative projects such as the ENCODE (76), FANTOM (77) and Roadmap Epigenomics consortia (1). However, the goal of our work was not to define the ultimate set of enhancers, but rather to verify if accounting for chromatin 3D organization can improve ETG pairing.

Thus, we primarily relied on Roadmap Epigenomics dataset as i) it covers a broad range of cell and tissue types; ii) it adopted shared protocols and quality standards, which is preferable to merging data from heterogenous sources; iii) the use of enhancers defined with epigenomics data facilitates the comparison against previously published algorithms for ETG pairing. More specifically, we used the peaks called by Roadmap Epigenomics for DNase-seq and H3K27ac ChIP-seq to define active enhancer regions for each of the selected 44 cell and tissue types (see Materials and Methods section and Table S2A). The average number of cell-specific enhancers is 33,560 (Figure 2A), with average size 316 bp (Figure S1B). Their pairwise comparison showed on average 50.6% of similarity (JI on overlap) (Figure 2B and Table S2B). On the other hand, the mean JI for coverage was 24.4% (Table S2A), due to the variable range of enhancer region sizes. To define a comprehensive reference enhancer catalogue, we considered the union of genomic intervals for cell-specific enhancers, resulting in *n* = 347,777 enhancer regions, with average size 416 bp (Figure S1B), *i*.*e*., slightly higher than the cell-specific enhancers, as the final catalogue is derived from their union.

**Figure 2.**
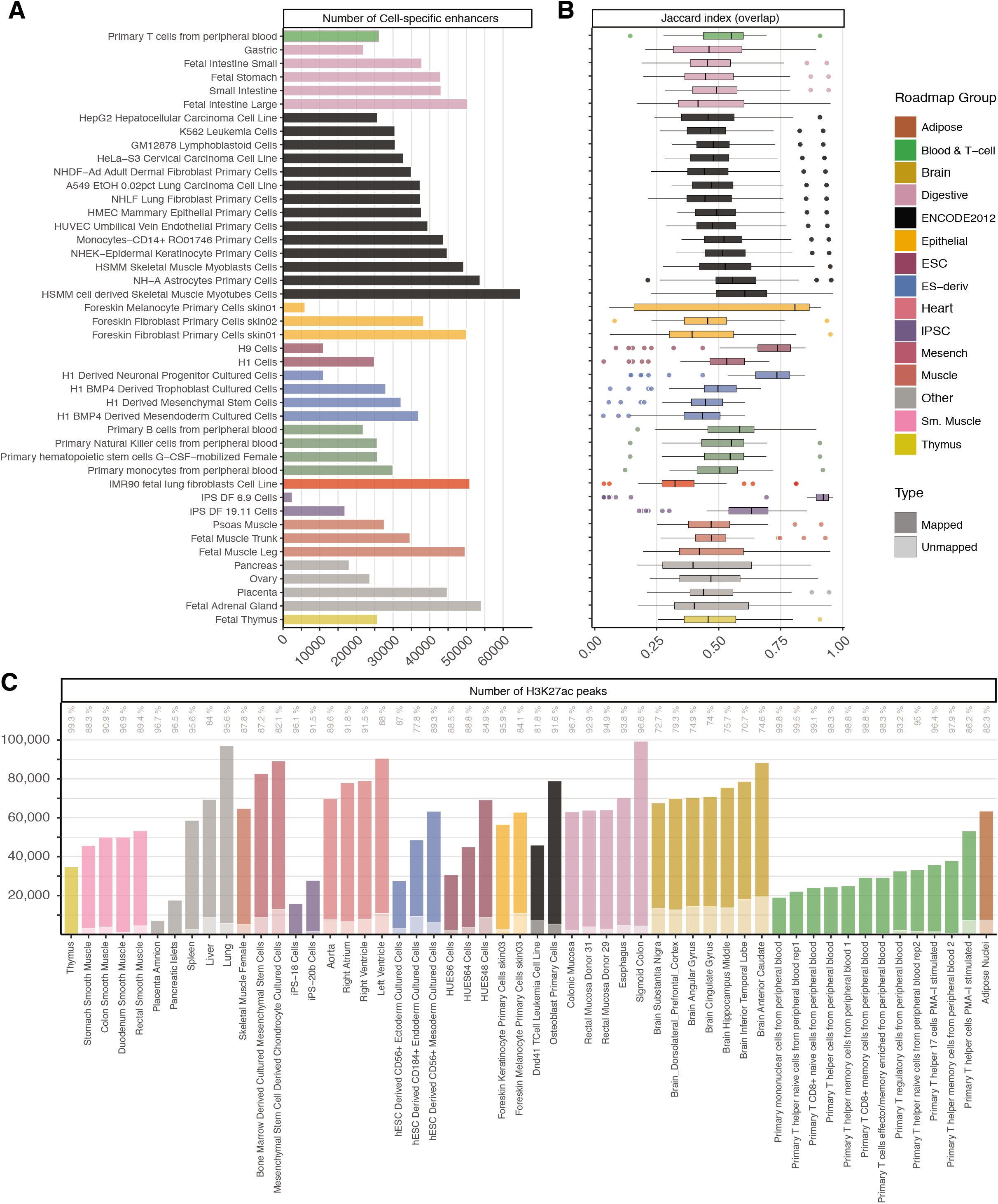
Definition of the reference enhancer catalogue. **A**. Number of cell-specific enhancers resulted by the intersection of DNase-seq and H3K27ac ChIP-seq peaks in a selected set of 44 cell and tissue types collected by the Roadmap Epigenomics consortia, coloured by Roadmap groups. **B**. Similarity (Jaccard index on overlap) among cell-specific set of enhancers. Each data point in the boxplots represents the ratio between the intersection of two cell-specific sets of enhancers over their union, taking as reference the group on the row. **C**. Number of H3K27ac ChIP-seq peaks of 54 additional cell and tissue types from the Roadmap Epigenomics project that overlap (dark colour) or do not overlap (light colour) with the union of H3K27ac ChIP-seq merged peaks of the set of cell and tissue types used to define the reference catalogue of active enhancers. The percentages of total H3K27ac peaks extension overlap are reported in light grey colour.

This reference enhancer catalogue can be considered exhaustive and representative also for other cell types. To this concern, we considered 54 additional Roadmap Epigenomics H3K27ac profiles, that were omitted from our enhancer catalogue definition because they lack a corresponding DNase-seq profile (Table S2C). 91.3% of the additional cell-specific H3K27ac peaks overlap to the union of H3K27ac peaks across the 44 cell types considered above. This overlap is 89.9% if we consider its extension over the total in the respective cell type (Figure 2C). The large overlap can be considered indicative for the completeness of our catalogue.

We also compared our reference enhancer catalogue to enhancer definitions by CAGE, from the fifth release of the FANTOM project (Functional ANnoTation Of the Mammalian genome) (77). We found that 57% out of the 58,200 filtered FANTOM enhancers (median length 270 bp) were also represented in our catalogue (Table S2D). It is noteworthy that FANTOM enhancer definitions were based on functional data from a much larger set of cell and tissue types including 432 primary cells, 135 tissue types and 241 cell lines (808 in total) (38). Thus, we deem our strategy a good compromise as FANTOM is based on 18 times more cell and tissue types.

Finally, we compared our catalogue to an *in vivo* validated set of enhancers coming from the VISTA Enhancer Browser database (40). Out of the starting 894 filtered VISTA database enhancers (median length 1,676 bp) (Figure S1C and Table S2D), 55% are present in our enhancer catalogue. It is worth remarking that VISTA is made of enhancers validated to be active in mouse embryos at development day 11.5. Thus, it is based on a different model organism and a very specific embryonic development stage, as opposed to our epigenomics datasets, which are derived from human samples, including several from differentiated tissues and cells from adult individuals. Moreover, superimposing the filtered FANTOM5 enhancers, only a minor residual number is detected in addition to our enhancer catalogue (Figure S1C and Table S2D). This observation confirms that alternative functional genomics definitions of enhancers, such as the CAGE-based FANTOM5, are overall comparable to ours.

### Enhancer-Promoter interactions in the 3D context

Enhancer-promoter contacts are generally confined within the boundaries of TADs (78), *i*.*e*. structurally separated domains relatively insulated from surrounding regions. TAD boundaries are mostly stable across cell types, but their insulation is far from absolute. Moreover, it is possible to identify a hierarchy of TADs, as any given Hi-C contact matrix can be analysed at different scales to derive alternative definitions of insulated domains (33, 79).

In order to account for these known features of chromatin 3D organization, we devised the HCL score, which is proportional to the likelihood of 3D co-localization of EP pairs (see Materials and Methods) (Figure 1B). We used 11 high-coverage Hi-C datasets (on average 660 million aligned reads), covering 10 different cell and tissue types (Table S1B) and binned at 10 kb resolution. We then applied the LSD algorithm to identify TAD boundaries at multiple scales, thus obtaining different segmentations of the genome that account for the hierarchy of structural domains. The number and size of TADs show a trend related to the LSD DI-window size parameter. Namely, we find fewer and larger TADs when increasing DI-window (hierarchy level) (Figure S2A): with average number ranging from 18,254 to 8,953, and average size from 183 kb to 525 kb (Table S3A). This pattern is comparable across datasets, despite differences related to sequencing depth. The pairwise comparison of domain boundaries across datasets, and across hierarchy levels, shows an average JI (coverage) ranging from 44.9 (for DI-window 5) to 35.8 (for DI-window 50) (Figure S2B). These results are in line with previous studies (80) and with the notion that several TADs are conserved across cell types.

We mapped our catalogue of enhancers (347,777) and reference set of promoters (18,027) to the inferred TADs (Table S3B), which are expected to compartmentalize the interactions between distal regulatory elements and target genes, and then we computed the HCL score for each EP pair. About 70% of EP pairs are within the same TAD in 2 or more Hi-C datasets, with a frequency distribution that is similar across all TAD hierarchy levels (Figure 3A). Nevertheless, the number of EP pairs grows as the hierarchy considered increases, as expected.

**Figure 3.**
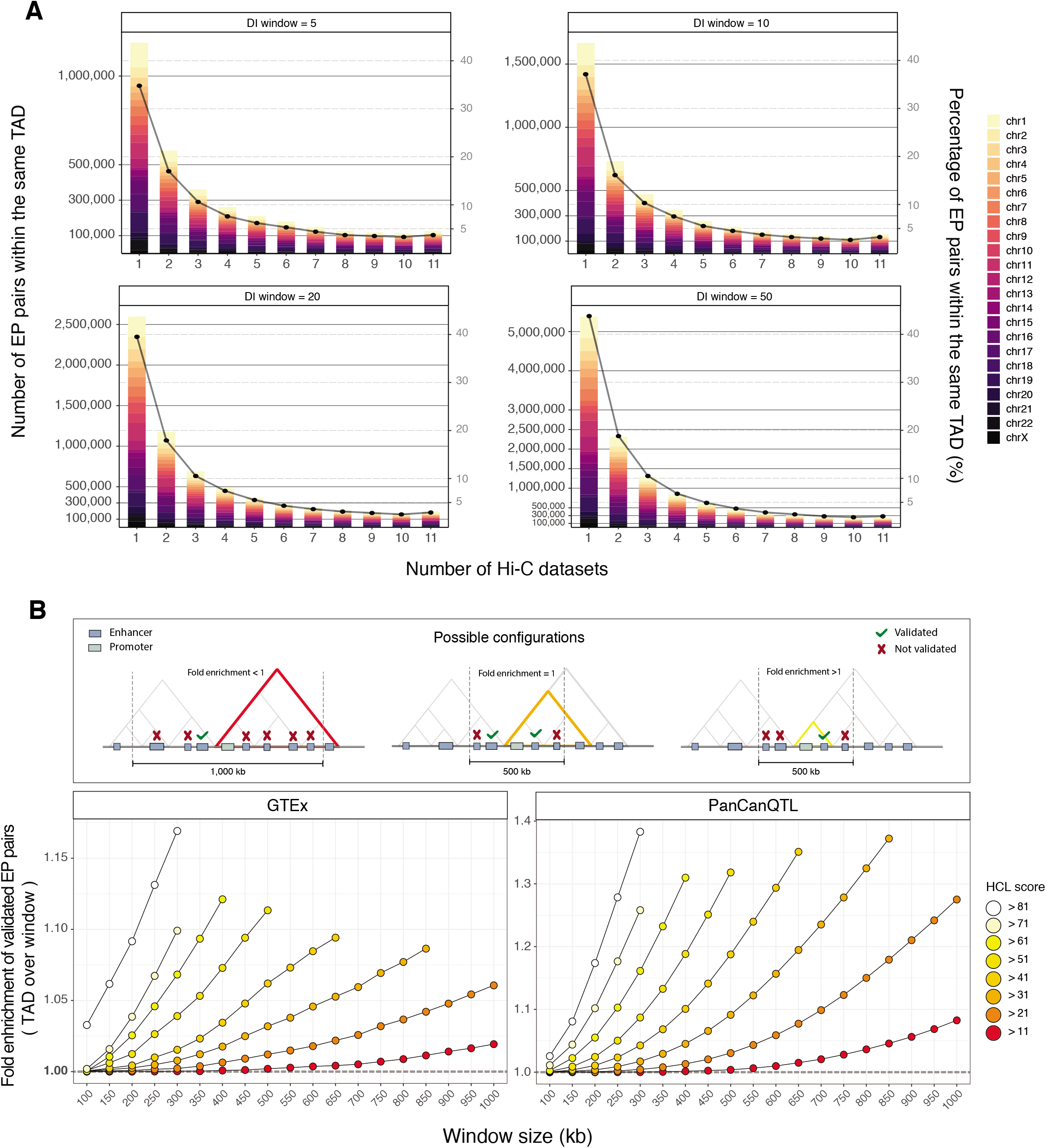
Enhancer-promoter interactions in the 3D context. **A**. Number (bar, left y-axis) and percentage (point, right y-axis) of EP pairs located within the same TAD for one or multiple of eleven analysed Hi-C datasets (x-axis), considering different TADs hierarchical levels (*i*.*e*., DI-window, panels), grouped by chromosomes (colours). **B**. Fold enrichment (y-axis) of eQTL-supported ETG pairs coming from two public datasets (GTEx, left panel; PanCanQTL, right panel) with respect to candidate pairs as defined with HCL score over a fixed-width window around the promoter of the target gene, for different cut-offs (score, colours; window-width, x-axis). As illustrated in the cartoon on top of the figure, values greater than one imply an enrichment of TAD-based pairing over results obtained with a fixed-window.

The EP pairs mapped within at least one TAD definition (i.e., HCL score > 0) are 12,949,150 (Table S3C) of which approximately 75% have a weak score (HCL score ≤ 11), *i*.*e*., they are supported only in few combinations of datasets and hierarchy levels. Possible scenarios above this threshold include EP pairs supported by all 11 datasets in at least one hierarchy level, or EP pairs supported by 2 or more datasets across multiple hierarchy levels. We deliberately designed the score to give comparable weight to these alternative situations.

This pattern, together with the overall trend observed in the HCL score, is robust respect to the TADs calling algorithm that is adopted. For example, we used as alternative input the TADs called by TopDom (52), another method that allows calling TADs at different levels of resolution by adjusting a tuning parameter (see Materials and Methods). We observed a similar percentage of poorly-supported pairs (79.8% with HCL score ≤ 11), as well as a per chromosome average 0.76 correlation (Spearman) between HCL scores based on TopDom or LSD TADs (Table S3D). While our observations are robust to different TADs calling algorithms, we noted that the number of candidate EP pairs based on LSD TADs is almost totally (98%) included in the set based on TopDom TAD definitions (Figure S2C). To this concern, it should be considered that the total number of candidate EP pairs for TopDom is approximately three-fold higher (38,527,013 pairs). This disparity is partly explained by the variable TAD calling performance of the two algorithms across datasets, which tend to give more different results when the coverage is lower (Figure S2D). Nonetheless, the comparison and benchmarking of TAD callers is beyond the scope of this work and has been extensively addressed in previous literature (19, 81, 82).

To ensure the robustness of the downstream results, we discarded poorly-supported EP pairs (HCL score ≤ 11) from subsequent analyses, as they may be the consequence of noise in the data depending on technical variables (e.g. coverage). This filter resulted in a total of 3,192,806 candidate ETG pairs.

There is a consensus on the fact that enhancers may not target the closest gene, in terms of linear sequence of the genome. Nevertheless, previous literature on genome-wide reconstruction of ETG pairs often adopted a fixed-width window around TSS to restrict the EP pairs search space. A commonly adopted boundary is a 1 Mb window (+/− 500bp around the TSS) to define the initial set of candidate pairs (83). Likewise, also the literature on expression quantitative trait loci (eQTLs) adopted a similar simplification. Indeed, cis-eQTL are often defined as SNPs within a 1 Mb window around the TSS, as opposed to trans-eQTL if the SNP falls beyond that distance threshold (or beyond 5 Mb for some other studies) or in another chromosome (84).

To verify if the use of chromatin 3D architecture as incorporated in the HCL score brought an advantage over the standard choice of a fixed-width window, we used a true positive set of ETG pairs based on eQTLs from the GTEx project (60) and PanCanQTL (61). Specifically, we verified the proportion of eQTL-supported ETG pairs with respect to the total number of considered pairs as defined with HCL or fixed-width windows. Since eQTLs are explored only for SNPs at a maximum distance of 1 Mb from the candidate target genes, to make a fair comparison we removed from our list all candidate pairs more distant than this threshold for a total of 3,102,154 remaining candidates. This filter was applied also for any subsequent analysis in which eQTLs were considered for comparison. We observed that in both eQTLs datasets there is generally a higher frequency of validated pairs when accounting for the chromatin 3D architecture, even if varying the threshold on HCL score and fixed-width window parameters (Figure 3B). These results highlight the existence of a stronger relationship between eQTLs and 3D distance, rather than linear distance.

### Physical proximity increases power of detection

We then reconstructed the enhancer regulatory network by integrating information on physical co-localization of EP pairs (HCL) and their activity synchronisation (CCA).

Enhancers are expected to show the properties of an active regulatory region in the specific cell context where they are contributing to activate a target gene (85). Thus, searching for “synchronised” enhancer and promoter activity across multiple cell types is a commonly adopted strategy in ETG pairing literature (86), although there is no consensus regarding a measure that best conveys their synchronisation. Differently from previously published methods, we adopted CCA as a convenient statistical framework to assess the synchronisation between activity of enhancers and promoters across multiple cell types in a fast and efficient way. We chose CCA because i) it is flexible with regard to the set of input functional genomics data used to estimate the activity level of enhancers and promoters; and ii) it accounts for the confounding factor of multiple types of functional genomics data being correlated with each other (Figure 1A).

In our case we used a combination of DNase-seq and ChIP-seq enrichment profiles to quantify the activity of enhancers and promoters. Namely, we used the maximum enrichment of DNase-seq and H3K27ac ChIP-seq for enhancers (347,777) and DNase-seq, H3K27ac and H3K4me3 for the promoters (18,027), as described in *Material and Methods* (Figure S3A). To minimize the influence of possible outliers and make the distributions comparable across all cell types, we used log_2_(*x* + 1) transformed enrichment values and adopted a chromosome-wise quantile-normalisation, respectively. We also tested cycle loess and variance stabilizing normalisation (VSN) as alternatives. They all yield similar results (Figure S3B-C), thus we selected the quantile normalisation as it preserves the original range of values.

To assess the association between the i-th enhancer and the j-th promoter, we performed the CCA considering the enrichment in these chromatin marks across the selected set of 44 cell and tissue types. The procedure retunrs a single p-value *p*^(*ij*)^, for each EP pair under consideration (see Materials and Methods). We performed CCA on the subset of EP candidate pairs filtered by HCL score >11, resulting in a total of *N* = 3,192,806 hypotheses to be tested (Table S3C). For each chromosome we estimated an adaptive L-value rejection threshold using the AdaPT procedure (58) with side information derived from the physical proximity of enhancer-promoter (HCL score)(Figure 1C). A representative example of the estimated thresholding rules for chromosome 19 is depicted in Figure 4A. We can see an increasing trend of the rejection curve in relation to HCL, implying that on equal nominal *p*-value our framework uses a less conservative significance criterion for the EP pairs showing higher likelihood of 3D contact interaction across cell types. Similar trends are observed for all chromosomes.

**Figure 4.**
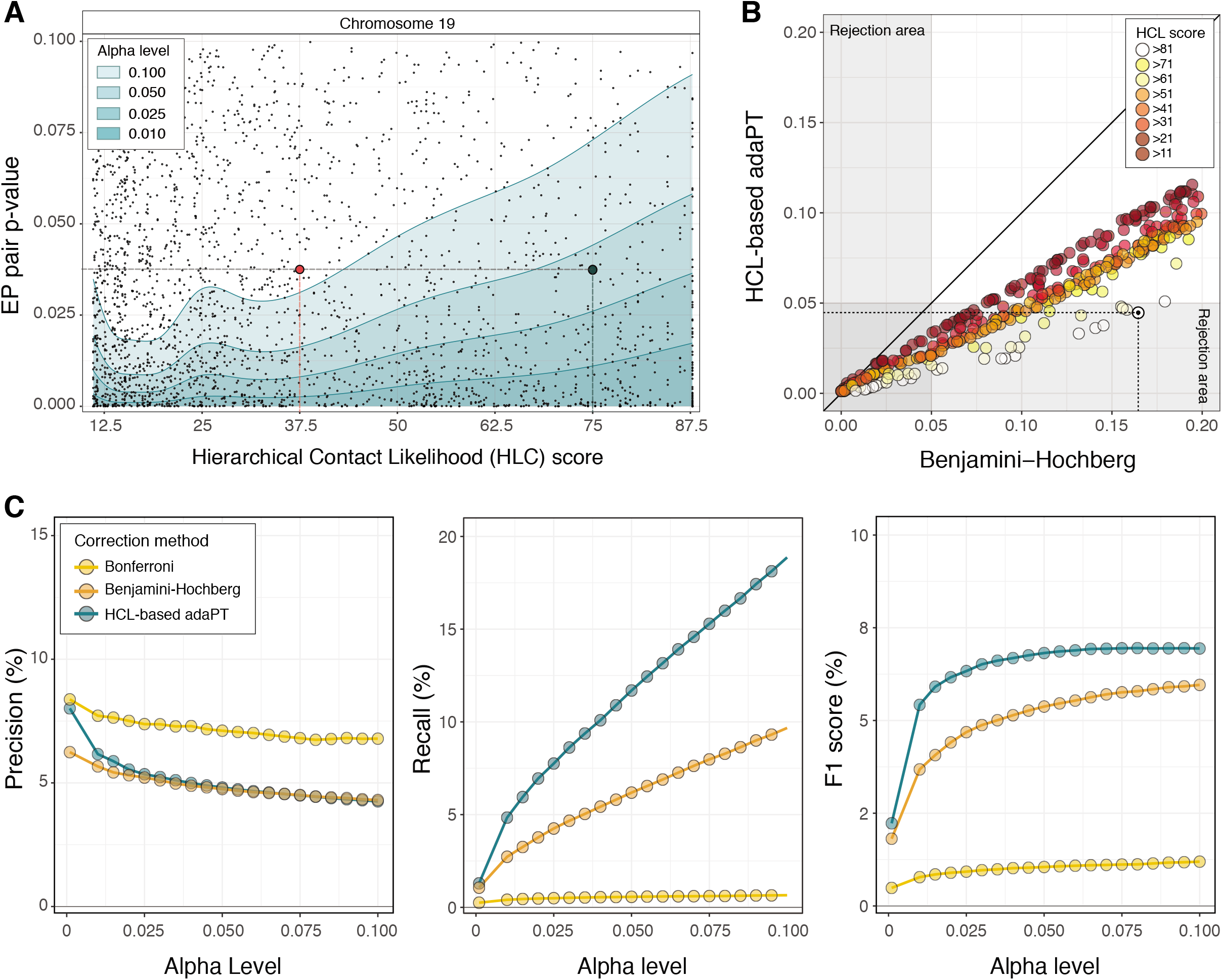
Physical proximity increases power of detection. **A**. Estimated iterative adaptive rejection threshold (AdaPT) leveraging side information derived from HCL scores (x-axis), for a subset of chromosome 19 at different alpha levels (green-shadow areas). Each point reports the p-value for the synchronised activity between an EP pair. Highlighted, two EP pairs with the same nominal p-values, but different HCL scores, for which the null hypothesis of independence is rejected at a confidence level of 0.95 (green point, high score), and not rejected (red point, low score). **B**. P-values associated to a subset of EP pairs for chromosome 19 adjusted with Benjamini-Hochberg (x-axis) and AdaPT (y-axis) approaches, coloured by HCL score classes. Highlighted with a solid black dot, an example EP pair located within the same TADs for all cell and tissue types and hierarchies (*i*.*e*., score>81), for which the null hypothesis of independence is rejected at a confidence level of 0.95 only by HCL-based AdaPT approach. **C**. Precision, recall and F1 score of predicted EP pairs based on eQTLs-supported ETG pairs (GTEX and PanCanQTL) over different alpha levels (x-axis), adopting three different multiple-testing correction approaches: Bonferroni (yellow), Benjamini-Hochberg (orange) and HCL-based AdaPT (green).

To illustrate the improvement achieved by integrating the score on chromatin 3D architecture, we adjusted the CCA *p*-values using Benjamini-Hochberg (BH) and HCL-based AdaPT correction (Figure 4B), and Bonferroni approach. Using the union of the GTEx and PanCanQTL datasets as reference true positive benchmark, we assessed the three methods in terms of Precision, Recall and F1 score (Table S4A). In Figure 4C we observe that, regardless of the level of confidence chosen, integrating the HCL in the p-value correction leads to an appreciable increase of the power (almost twice) without affecting the accuracy of the predictions.

The complete list of candidate enhancer-promoter pairs annotated with the HCL score, corrected and uncorrected p-values, validations according to multiple reference datasets are publicly released (see Data Availability). For the subsequent analyses we maintained the EP pairs with HCL-based AdaPT adjusted p-value ≤ 0.05 resulting in a total of 233,304 predicted pairs.

### Benchmarking against other ETG pairing methods

To benchmark our ETG pairing framework against other methods, as described more in details in the Materials and Methods section, we selected 7 algorithms (Table S1G): FOCS, PreSTIGE, RIPPLE, PETmodule, TargetFinder, JEME and DeepTACT (71) (72) (73) (22) (23) (21)(74). Overall, these approaches represent the evolution of ETG predictors proposed between 2014 and 2019, covering three of the main categories as defined by Hariprakash and Ferrari (75) (i.e., correlation, supervised learning and regression-based methods).

It is worth noting that previous publications used different definitions for the reference set of enhancers and promoters, thus introducing heterogeneity in the input candidate pairs which we considered in our analyses.

The ETG pairs called by the selected tools showed prominent differences in terms of EP distance distributions (Figure S4A). For this reason, in addition to the eQTLs datasets we also considered as true positives a set of 9 capture Hi-C (cHi-C) datasets (62–68) (Table S1E), designed to identify contacts between promoters and distal chromatin regions. Taken together, the two types of data provide a broader coverage of functional and physical interactions occurring at different distance ranges. Namely, eQTLs and cHi-C data are representative of mid-range (average distance of 82 Kb) and long-range (average distance of about 326 Kb) interactions, respectively (Figure S4B).

Figure 5 and Table S4B summarize the performance of algorithms grouped into two categories: *i*.*e*. methods that rely on i) a unique set of promoters and enhancers as input (FOCS and our framework) or ii) cell-type specific definitions of enhancers and promoters (PETmodule, JEME, PreSTIGE, RIPPLE, DeepTACT and TargetFinder). In the first group performances are assessed directly on the unique list of ETG pairs, whereas in the second group performances are reported as average across the cell-type specific lists.

**Figure 5.**
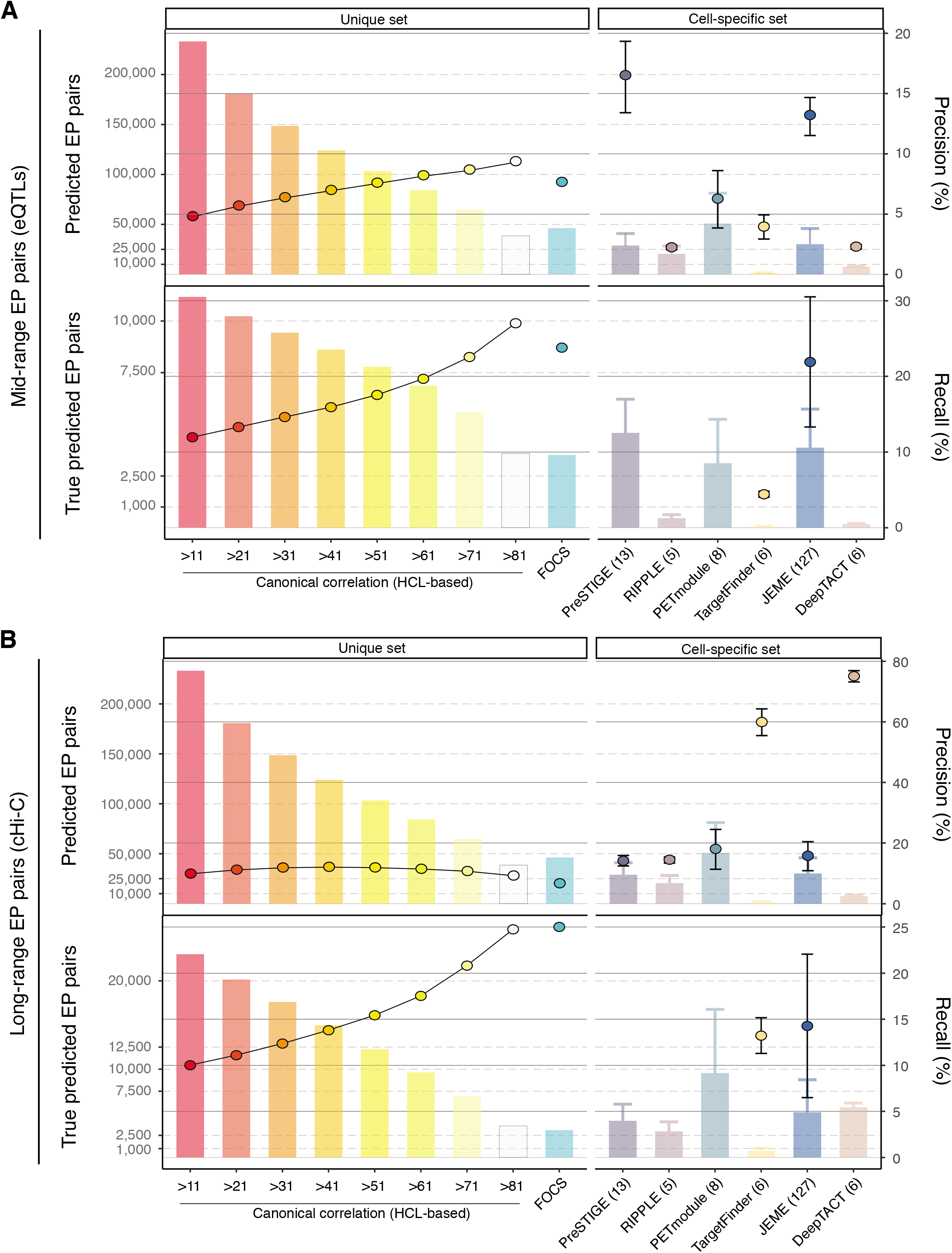
Benchmarking against other ETG pairing methods. Performances of our approach (evaluated at different HCL score cut-offs) and other seven ETG pairing algorithms assessed based on mid-range (**A**, eQTLs supported) and long-range (**B**, cHi-C supported) true positive EP interactions. Bars (left y-axis) report the number of predicted (upper panel), and true predicted EP pairs (bottom panel). Points (right y-axis) report precision (upper panel) and recall (bottom panel). Recall is available only for tools for which the list of EP candidate pairs were released. Algorithms are grouped in two categories: methods that rely on a unique set of promoters and enhancers as input (left panels) or rely on cell-type specific definitions of these sets (right panels). For methods in this last category, the standard deviation (whiskers) and the number of evaluated cell and tissue types (text in brackets) are reported.

Considering mid-range interactions (based on eQTLs, Figure 5A), our method ranked above most of the other algorithms with a precision ranging from 4.8% to 9.4%, depending on the HCL score cut-off. Only JEME and PreSTIGE showed remarkable performances with 13.1% and 16.4% precision, respectively. However, our method exhibited recall values ranging from 12% to 27%, depending on the HCL score cut-off, which were comparable to JEME (21.9%). The recall cannot be computed for PreSTIGE and other methods not providing the starting set of candidate ETG pairs.

Instead, focusing on FOCS, which among the selected algorithms is the only other one with a unique EP list, our approach showed better performances for HCL scores greater than 71, while a decline was observed for the remaining cut-offs. This pattern is mainly due to the use of a candidate EP pairs which is about 16 times larger than FOCS (3,099,004 versus 192,800 pairs) and the exploration of interactions over longer distances, up to 8 times more distant (average distance: 334 kb versus 42 kb). These two peculiarities result in a large imbalance in the initial proportion of true pairs, making their detection more challenging. Indeed, considering an index that is not affected by this bias (i.e., the relative improvement, RI), we estimated that the observed to expected ratio of true pairs (Table S4B) in FOCS was equal to the random choice over the initial candidate pairs (0.99), contrary to our algorithm (from 1.15 to 1.59).

Instead, when considering long-range interactions (based on cHi-C, Figure 5B), we observed the highest precisions in DeepTACT (75%) and TargetFinder (60%), whereas all the other methods are comparable, with values around 13%. Although the precision for our approach was slightly lower than the other tools (ranging from 9.2% to 12.1% with different HCL thresholds), the recall proved to be good (ranging from 10% to 24.5%). Interestingly, the better-performing algorithms in the long-range interactions coincide with the worst-performing ones in the mid-range interactions. Conversely, our approach has good performances in both conditions.

### Cell specificity of the predicted enhancer-promoter pairs

To further appraise the performance of our method, and investigate its behaviour in identifying cell type-specific ETG pairs, we performed an additional direct comparison with JEME. This choice was motivated by the consideration that JEME is the most comprehensive in terms of cell-type specific ETG lists (127 cell types from the Roadmap Epigenomics dataset, Table S1G) and overall resulted as the best-performing among the 7 algorithms selected for our benchmarking.

We used JEME initial set of enhancers, genes and candidate ETG pairs. This choice allows us to minimise the sources of heterogeneity in the comparison, and to test the flexibility of our algorithm with inputs other than those used as reference in our work. In particular, we considered all 127 cell-type specific lists of candidate ETG pairs provided by JEME authors. It must be noted that JEME does not return any threshold for FDR control. Thus, we compared the methods by calculating only the precision based on the top-ranked ETG pairs using different cut-offs (1000, 2000, 3000, 4000 and 5000). As reported in Figure 6A and Table S4C, the overall precision of our method (ranging from 16.6% to 19.7%) is comparable to JEME (from 19.8% to 20.1%). However, the precision for the long-range interactions is slightly higher in JEME (from 10.2% to 11.6%) as opposed to our method (from 6.5% to 9.9). We also noted a large variance in JEME estimates and an increasing precision in our method with less stringent cut-offs. It is worth remarking that our choice of using JEME definitions of enhancers, genes and candidate ETG pairs, might put JEME in an advantageous position and render the comparison of the two approaches somewhat biased.

**Figure 6.**
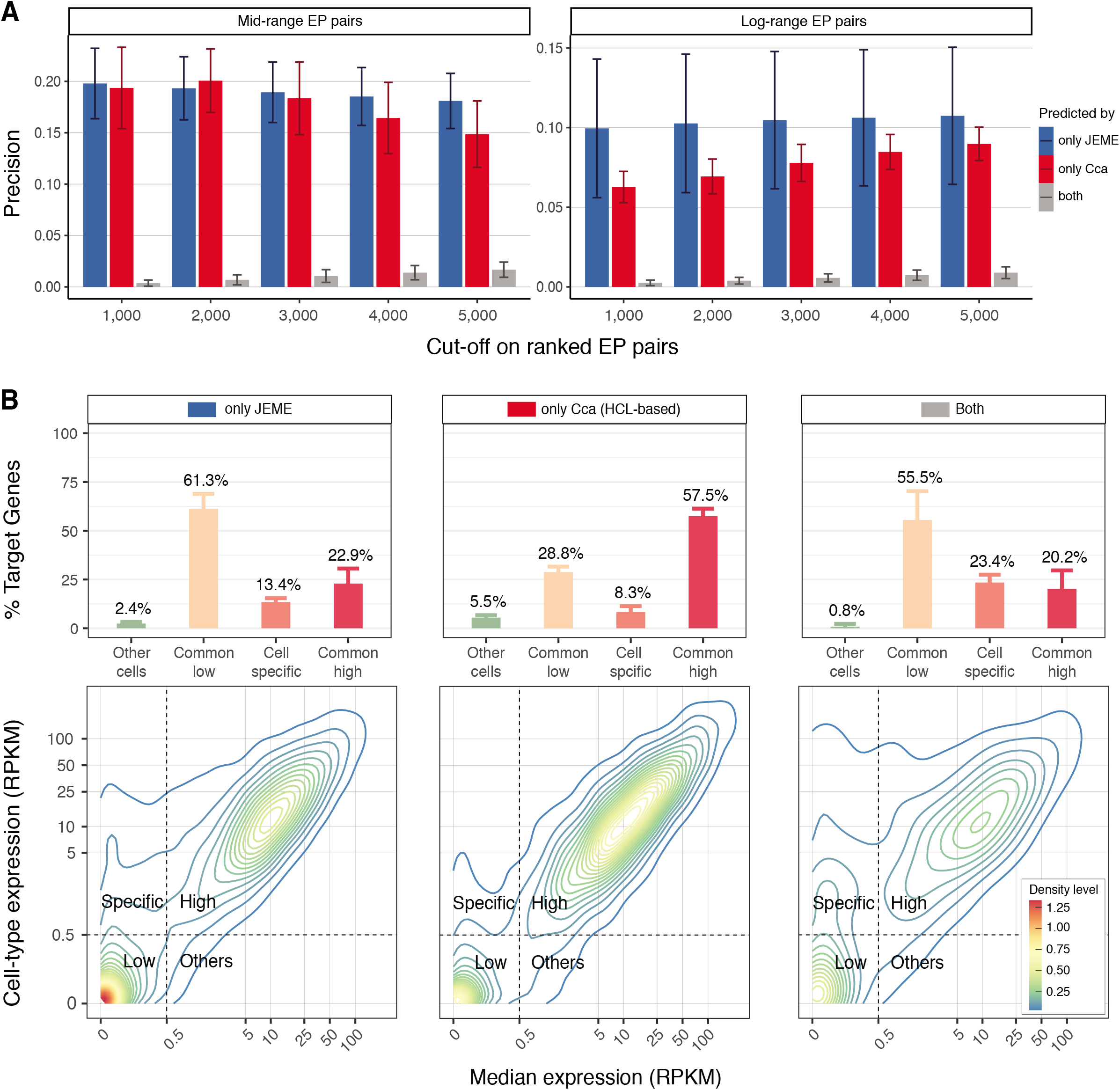
Cell specificity of the predicted enhancer-promoter pairs. **A**. Average precision (bars) and standard deviation (whiskers) assessed based on mid-range (eQTLs supported, left panel) and long-range (cHi-C supported, right panel) true positive interactions for ranked predicted ETG pairs by only JEME (blue bars), only our framework (red bars) and both tools (grey bars) in 127 cell types coming from Roadmap Epigenomics datasets for different cuts-off of the ranked lists of predictions. Initial sets of enhancers, genes and candidate ETG pairs are the ones described in the original publication of JEME. **B**. Gene expression density (contouring plots, bottom panels) and percentages (bars, upper panels) of target genes on top 1,000 predicted interactions by only JEME (left panels), only our framework (middle panels) and both tools (right panels) in matched 57 out of 127 cell types considered in the original publication of JEME. Contouring plots are calculated merging the set of predicted target genes (x-axis: expression of the target gene in the cell type considered; y-axis: median expression in all the cell types) for each of the 57 cell types. For each cell type, a target gene is classified based on its expression in the specific cell type versus the overall expression profiles in all the cell types as: commonly low (salmon, common low) or highly expressed (dark pink, common high), expressed only in the cell type considered (light pink, cell specific) or expressed only in a small subgroup of other cell types (light green, other cells). The threshold used for the classification is highlighted with dotted grey lines in the contouring plots.

The most striking result is the almost complete lack of overlap between the ETG pairs identified by JEME and our approach: on average only 0.6% of the true pairs are identified by both approaches. To exclude the possibility that the discrepancy was due to a bias of our method, we used matched gene expression data on 57 out 127 considered cell types, to verify the predicted target genes activity. As reported in Figure 6B, although JEME identified a slightly higher percentage of targets that are strictly cell-type specific (13.4% versus 8.2%, respectively), more than a half are generally low expressed genes (61.3% versus 28.8%). On the contrary, our framework leads to the identification of targets with high expression (22.9% versus 57.5%), thus confirming its reliability and versatility in detecting relevant ETG pairs with results generalizable to other cell types.

### Benchmarking against independent reference dataset

As described above, the benchmarking against previously published algorithms is hampered by the extreme variability in solutions adopted to define the starting set of candidate ETG pairs. Therefore, we further compared our results against the only independent reference benchmarking dataset for ETG pairing currently available: BENGI (Benchmark of candidate Enhancer-Gene Interactions) (69). This database contains a collection of uniformly processed datasets that integrate the Registry of candidate cis-Regulatory elements (cCREs) (76) with experimentally derived genomic interactions (Table S1F). Thus, we used their enhancers, genes and candidate ETG pairs as input for our framework.

To appreciate the distinct features captured by the experimental datasets used to curate BENGI interactions, we computed precision-recall (PR) curves for the GM12878 cell line (a lymphoblastoid cell line), which is the most extensively surveyed one in BENGI (Figure 7A and Figure S5A). The PR curves include: AdaPT corrected p-values (sorted in ascending order); canonical correlation (decreasing order); HCL score (decreasing order); EP distance (increasing order). We noticed two scenarios: i) in top-ranked pairs supported by ChIA-PET or 3C-derived methods, which include the majority of the BENGI validated interactions (1,706,837 pairs, 95%), the distance does not provide any insight in the identification of true EP pairs, resulting in close to random selection or worse; ii) instead in eQTL datasets (87,982 pairs, 5%), the distance classifier remains consistently above the performance of our method, although CCA is initially aligned. It is also worth mentioning that a true positive EP pair in one BENGI experimental dataset list could be a true negative in another BENGI list.

**Figure 7.**
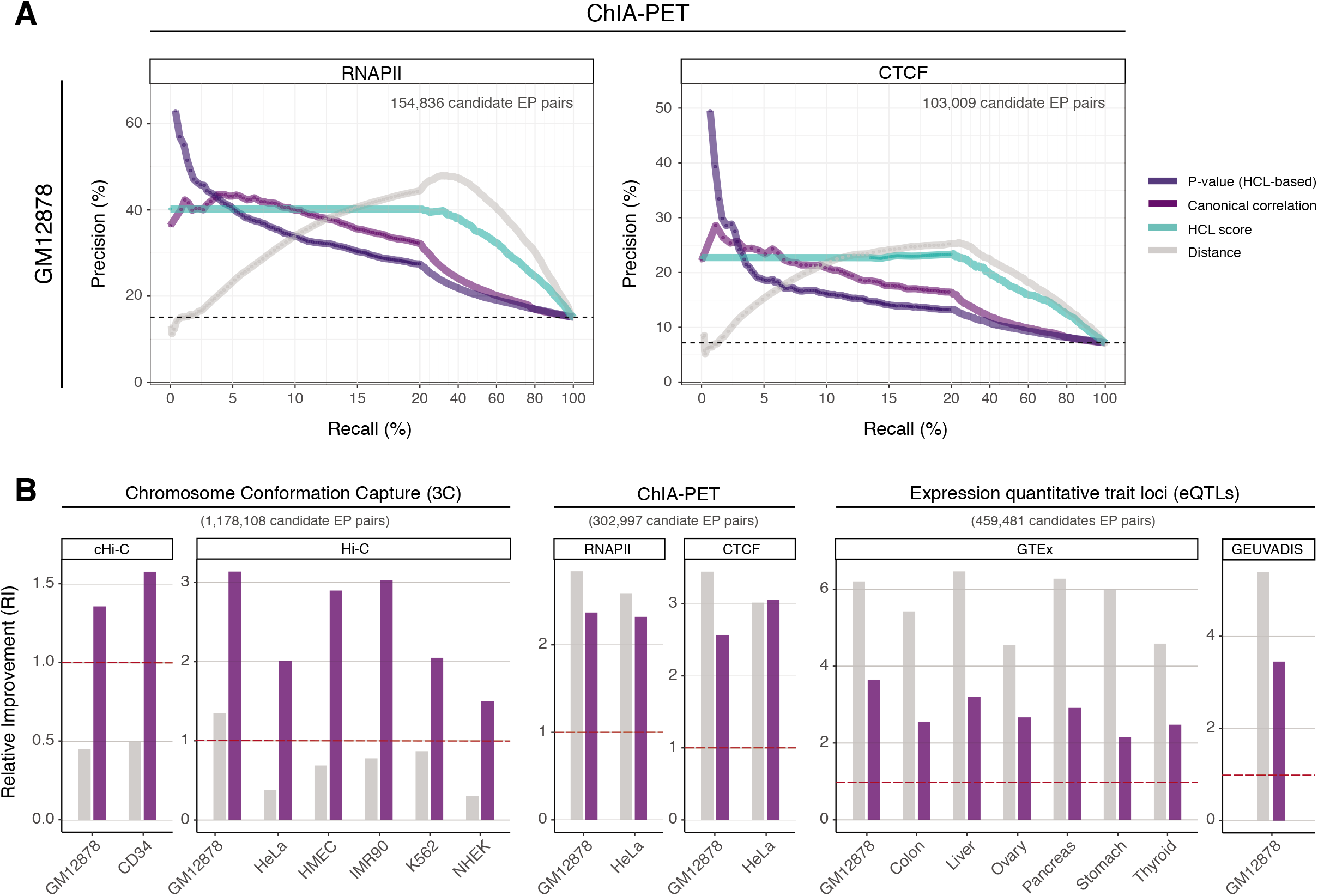
Benchmark against independent reference datasets. **A**. Precision-recall curves for GM12878 cell line BENGI benchmark datasets, assessed employing two datasets of ChIA-PET supported true ETG positive interactions (RNAPII, left panel; CTCF, right panel). Performances are calculated for: AdaPT HCL-based adjusted p-values (dark purple lines), canonical correlation (light purple lines), HCL score (aquamarine lines) and linear EP distance (grey lines). The total number of ETG pair considered is reported on the upper right corner of each panel. The plot is reporting an expanded x-axis in the initial part of the curve (corresponding to recalls up to 20%) to provide more details in the most informative part of the chart. For higher recall values, the precision-recall curves reported here tend to converge without further crossing each other. **B**. Relative improvement (y-axis) for all twenty BENGI benchmark interactions, assessed employing different sources of true ETG positive interactions. Namely, chromosome conformation capture (left panels), ChIA-PET (middle panels) and expression quantitative trait loci (right panels). Random choice (RI=1) is marked by red dashed lines. Performances are calculated for AdaPT HCL-based adjusted p-values (dark purple bars) and linear EP distance (grey bars), based on the same cuts-off on FDR=0.01 for adjusted p-values.

To further clarify this observation, we considered the twenty individual experimental datasets used by BENGI and we computed the relative improvement (RI) achieved by our method or EP distance alone (Figure 7B and Table S4D). As shown, the pattern described above is consistent across all validation datasets. Indeed, our method obtains robust and significantly enriched performances (i.e., higher than random classification) both for eQTLs (from 2.15 to 3.65) and ChIA-PET or 3C-derived interactions (from 1.36 to 3.14). Instead, the distance alone yields performances lower than what expected by chance in 7 out of 8 datasets of 3C-derived interactions.

## DISCUSSION

Here we present a new approach to refine the pairing of enhancers and target gene promoters. The three principles underlying our approach are: **i) the flexibility**, with respect to all input data; **ii) the use of prior-knowledge**, as we leverage 3D chromatin architecture to inform EP pairing; **iii) the robustness of the statistical framework**, as the method does not require arbitrary parameter tuning decisions and guarantees statistical control of the false discovery rate.

**The flexibility** of the method allows the end users to provide any preferred definition for the reference set of enhancers, genes, and functional genomics data used to quantify their activity. This versatility is primarily guaranteed by the use of CCA which provides the foundation for a very general statistical framework. Indeed, other correlation tests, as well as several parametric tests (including ANOVA, linear or multivariate regression, discriminant analysis, and chi-squared test), can be described as special cases of CCA, as demonstrated in literature (87).

Then the **prior-knowledge** on chromatin 3D organization has been carefully considered in defining the HCL score to quantify EP pairs physical proximity. In fact, we took into account the general consensus in literature that ETG pairs occur within TAD domains, and that alternative TAD definitions at multiple scales are concurrently present in the cell population. This biology-derived knowledge has been directly used to compute the HCL score integrated as side information in the adjustment of CCA *p*-values for each EP pair.

This may seem a counter-intuitive solution as opposed to directly using the EP loci contact frequency from the Hi-C matrix. However, using the multiscale TAD structure instead is a fundamental paradigm shift that allows overcoming relevant technical limitations. First of all, Hi-C contact matrices are generally binned at few kb resolution, thus at a scale that does not allow distinguishing regulatory regions close to each other. Indeed, even the most recent ETG pairing attempted with this strategy could not go beyond 5kb resolution (88). Moreover, Hi-C point interaction calling algorithms have been shown to yield very variable results even across biological replicates (19). Finally, Hi-C point interactions are generally considered to be very dynamic across cell types, thus they would always require a precise match between the cell and tissue type used for Hi-C and other genomics data required for the ETG pairing.

Instead, TADs are expected to be more conserved across cell types, thus allowing to extend the applicability of the HCL score to different cell contexts.

It is also worth remarking that our method is flexible for what concerns the hierarchy of TAD structural domains provided as input. As such, the end user may adopt the preferred algorithm for calling TADs at multiple scales.

Additionally, **the robustness** of the method is safeguarded by the use of AdaPT multiple testing correction which is combining the CCA p-value and the HCL score for each EP pair. This solution increases statistical power by prioritizing most promising hypotheses based on side information. As a representative example attesting the importance of such strategy, we noted the HBB-LCR region, which is a known complex distal regulatory region, responsible for the coordinated regulation during development of the human beta globin genes (HBE1, HBG2, HBG1, HBD and HBB). The LCR region contains several enhancers that are paired to one or multiple beta globin genes in our list of EP pairs (see Data Availability). For several of them, the BH adjusted p-value is not significant, whereas the AdaPT adjustment is able to detect a significant association to the beta globin genes. In particular, the enhancer at chr11:5297767-5298471 has a significant AdaPT p-value for all of the five beta globin gene promoters, whereas the same enhancer would be significantly associated only to HBG1 and HBG2 based on BH correction.

We also note that our strategy of using HCL as side information to control FDR is in principle a generalizable approach, that could also be applied to p-values for ETG pairs coming from other methods. Unfortunately, all the methods that we surveyed (70) did not really address the multiple testing correction problem, as most of them are either based on a classifier, or some other custom score, thus not returning an actual p-value for individual EP pairs.

Thus, we reconstructed the ETG regulatory network by applying our framework using genome-wide profiles of epigenetic marks for 44 cell and tissue types, together with multi-scale TAD calls derived from 11 high-coverage Hi-C datasets. To this concern, it may be worth remarking that we quantified the gene activity using the epigenetic marks at promoters, as opposed to the expression level of the gene transcripts. This is a commonly adopted choice in the literature of this field. The rationale behind this solution is based on the role of enhancers in triggering transcription: hence enhancers will show a synchronized activity with markers of transcription initiation in their targets. Instead, the actual transcripts abundance will depend also on multiple levels of co-transcriptional (e.g. RNA polymerase pausing, processivity in elongation, splicing and poly-adenylation) and post-transcriptional regulation (e.g. mRNA stability). All of these mechanisms will confound the synchronisation between enhancers activity and transcriptional output.

We identified a total of 233,304 EP pairs with adjusted p-value ≤ 0.05 and extensively benchmarked our results against 7 pre-existing algorithms representing multiple categories of ETG pairing methods. We used multiple sources of true positive ETG pairs, including eQTLs, cHi-C and a recently published curated database (BENGI), which is relying as well on multiple sources of experimental data.

We observed consistent performances in both mid- and long-range interactions for our method, as opposed to previously published algorithms that generally perform better on one of the two distance ranges (Figure 5). Moreover, we showed that our method compares well also to algorithms capturing cell-type specific ETG pairs (Figure 6), even though we aim to provide a generalizable ETG network that can be extended to multiple cell types.

It is worth noting that some of the previous algorithms based on supervised methods were actually trained using eQTLs or cHi-C data as true positive sets, that we have used for the benchmarking as well. Implying that other methods may have an advantage in the benchmarking statistics presented here.

We also must note that the definition of true positive ETG pairs may suffer some limitations. Namely, cHi-C and similar techniques can confirm a physical proximity between specific genomic regions, but this is not always resulting in a functional regulatory interaction between them. Likewise, eQTLs confirm a correlation between a gene expression and a genetic variants (SNP), but the actual distal regulatory region may be in a different position within the SNP linkage disequilibrium block.

Considering these limitations, which are anyway affecting any benchmark of ETG pairing algorithms, we further dissected our method performances on the BENGI database, containing an additional curated set of ETG pairs. Even in this case, our method confirmed consistent performances across all types of ETG supporting data, covering both mid- and long-range interactions (Figure 7B).

As discussed in detail by (89), performing a quantitative comparison of ETG pairing methods is a challenging task, where critical points should be considered such as i) properly separating training and validation sets; ii) considering the distance as a relevant feature affecting ETG pairing; iii) paying attention to different definitions of enhancer and promoter windows adopted by distinct algorithms. Throughout our work we carefully took into account these critical points as discussed in details for each individual analysis. The resulting framework proved to yield coherent results across different test datasets and cell types, thus confirming its value as a generalizable approach for ETG regulatory network reconstruction. In order to facilitate reproducibility of results, and widespread adoption in the community, we are publicly releasing the code and input datasets used for this study (see Data Availability). This tool will provide a valuable resource especially for translational studies aiming to annotate the functional role of non-coding sequence variants in distal regulatory elements. In particular, we envision possible applications in clinical genomics studies of cancer and undiagnosed genetic diseases.

## DATA AVAILABILITY

All public datasets used in this manuscript are described in Table S1.

The source code and the complete list of candidate enhancer-promoter pairs annotated with the HCL score, corrected and uncorrected p-values, validations according to multiple reference datasets are available at https://bitbucket.org/Elisetta88/3d-etg/.

## SUPPLEMENTARY DATA

Supplementary Data are available at NAR Online.

## ACKNOWLEDGEMENTS

We thank Chiara Romualdi for advice on statistical analysis and feedbacks. We thank Marco Morelli and Natoli Gioacchino for critical feedback on the manuscript. We thank Vincenzo Corbo and Pietro Delfino for stimulating discussions. We thank Cristiano Petrini for precious help with pipelines. We are grateful to Orso Maria Romano for support and constructive inputs on models and graphs.

## SUPPLEMENTARY FIGURE LEGENDS

**Figure S1.**
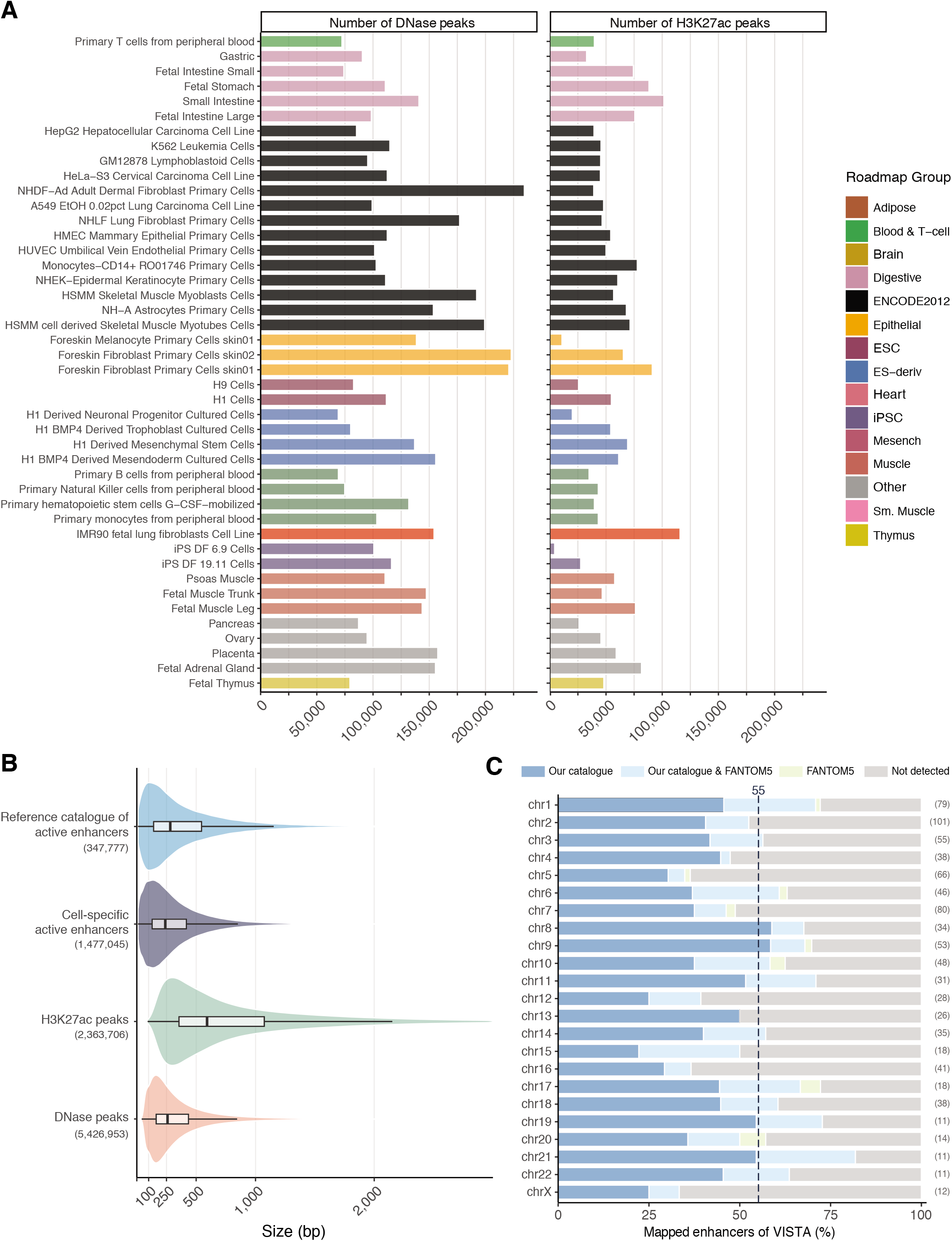
Definition of the reference enhancer catalogue. **A**. Number of DNase-seq and H3K27ac ChIP-seq peaks in a selected set of 44 cell and tissue types collected by the Roadmap Epigenomics consortium, coloured by Roadmap groups. **B**. Size distributions of H3K27ac ChIP-seq peaks, DNase-seq peaks, cell-specific active enhancer and active enhancers that belong to the reference catalogue (outliers not shown); the number of elements used for density estimation is reported in brackets. **C**. Proportion of *in* vivo validated set of enhancers coming from the VISTA Enhancer Browser database that overlaps only with our reference enhancer catalogue (blue), FANTOM5 set (yellow), both sets (light blue), otherwise (grey), grouped by chromosome. The number of considered *in-vivo* validated enhancers are reported in brackets.

**Figure S2.**
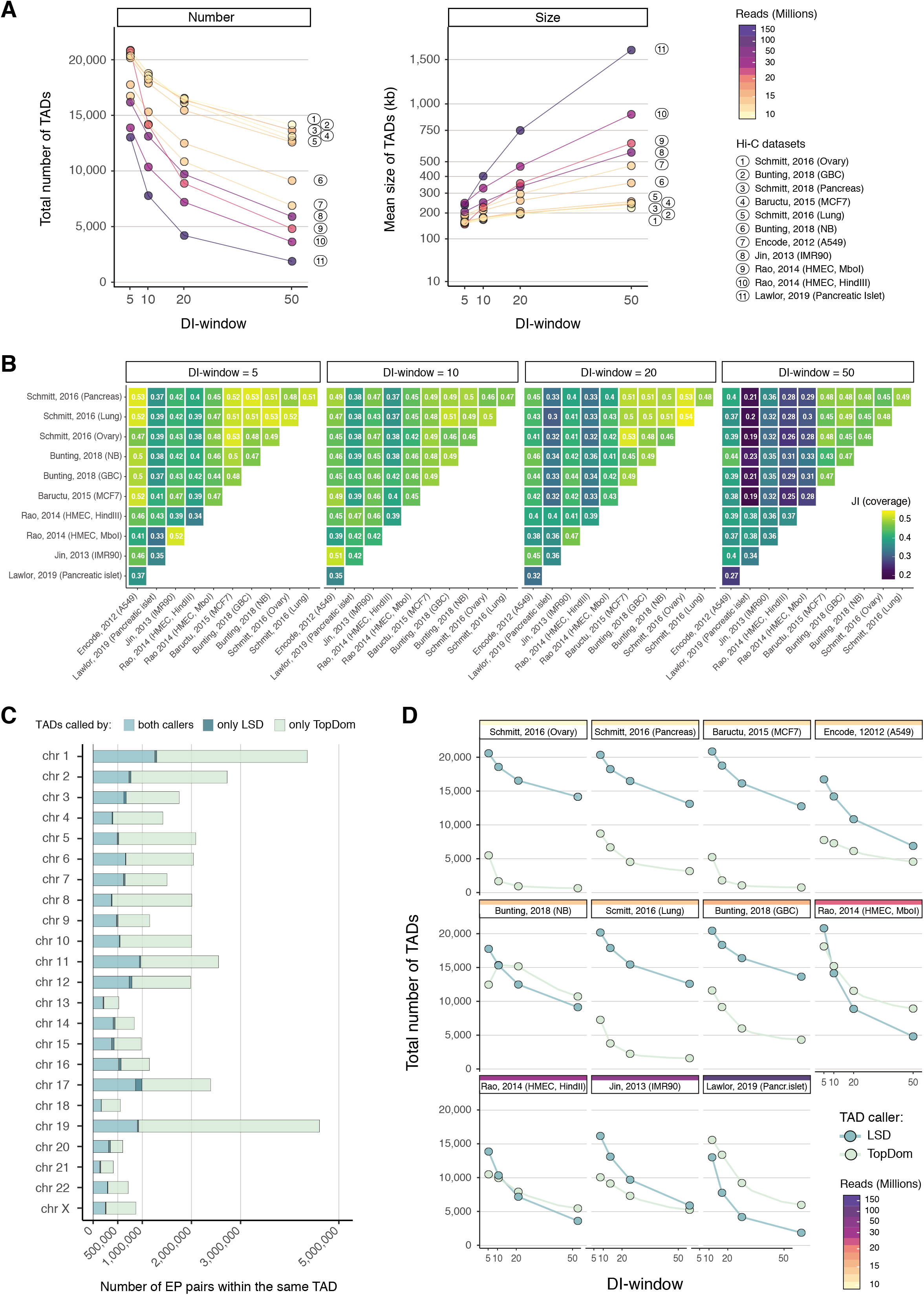
Enhancer-promoter interactions in the 3D context. **A**. Total number (left panel) and mean of the average size per chromosome (right panel) of TADs called in eleven Hi-C datasets, for decreasing hierarchical levels (*i*.*e*., DI-window parameters, x-axis). Hi-C datasets are coloured and sorted (ascending order) by sequencing depth (*i*.*e*., number of filtered reads). **B**. Pairwise Jaccard index on coverage of TADs, across hierarchical levels (panels) for the analysed eleven Hi-C datasets. **C**. Number of EP pairs located within the same domain boundaries only for LSD (dark green), TopDom (light green) or both TAD callers (middle intensity of green), grouped by chromosome. **D**. Total number of TADs called by LSD (dark green) and TopDom (light green) algorithms for decreasing hierarchical levels (*i*.*e*., DI-window or window parameters, respectively, x-axis) in eleven Hi-C datasets (panels). Hi-C datasets are coloured and sorted (ascending order) by sequencing depth (from upper-left to bottom-right panel).

**Figure S3.**
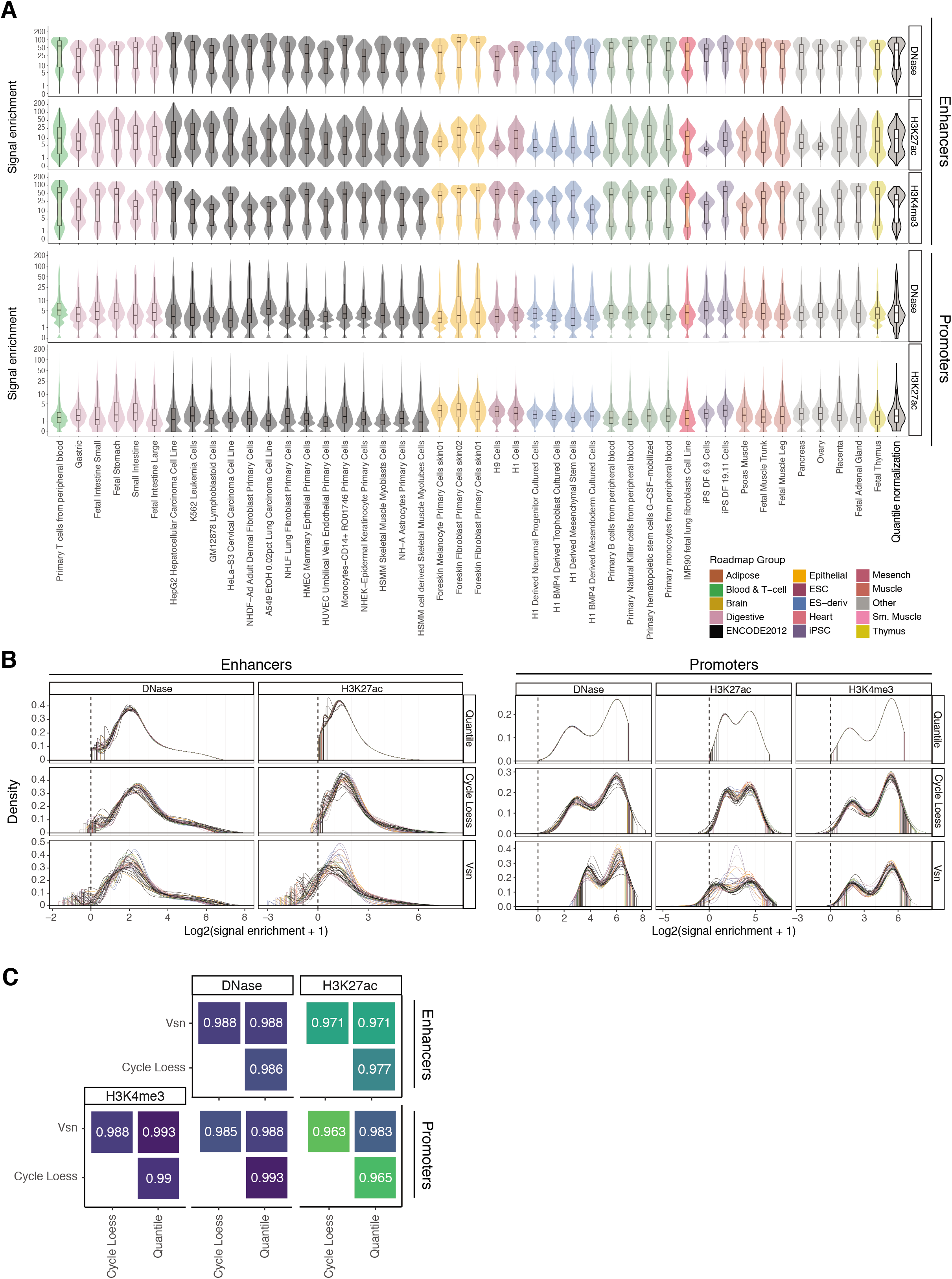
Physical proximity increases power of detection. **A** Representative example (chromosome 19) of signal activity distributions for the reference set of promoters (upper panels) and enhancers (bottom panels) in a selected set of 44 cell and tissue types collected by the Roadmap Epigenomics consortia, calculated as the maximum enrichment of DNase-seq, H3K27ac ChIP-seq profiles. Grey violin and boxplots report the signal activity distribution after chromosome-wise quantile normalization. **B**. Signal activity distributions in chromosome 19 for the reference set of enhancers (left panels) and promoters (right panels) after three different normalization strategies, and **C** their pairwise Spearman correlations.

**Figure S4.**
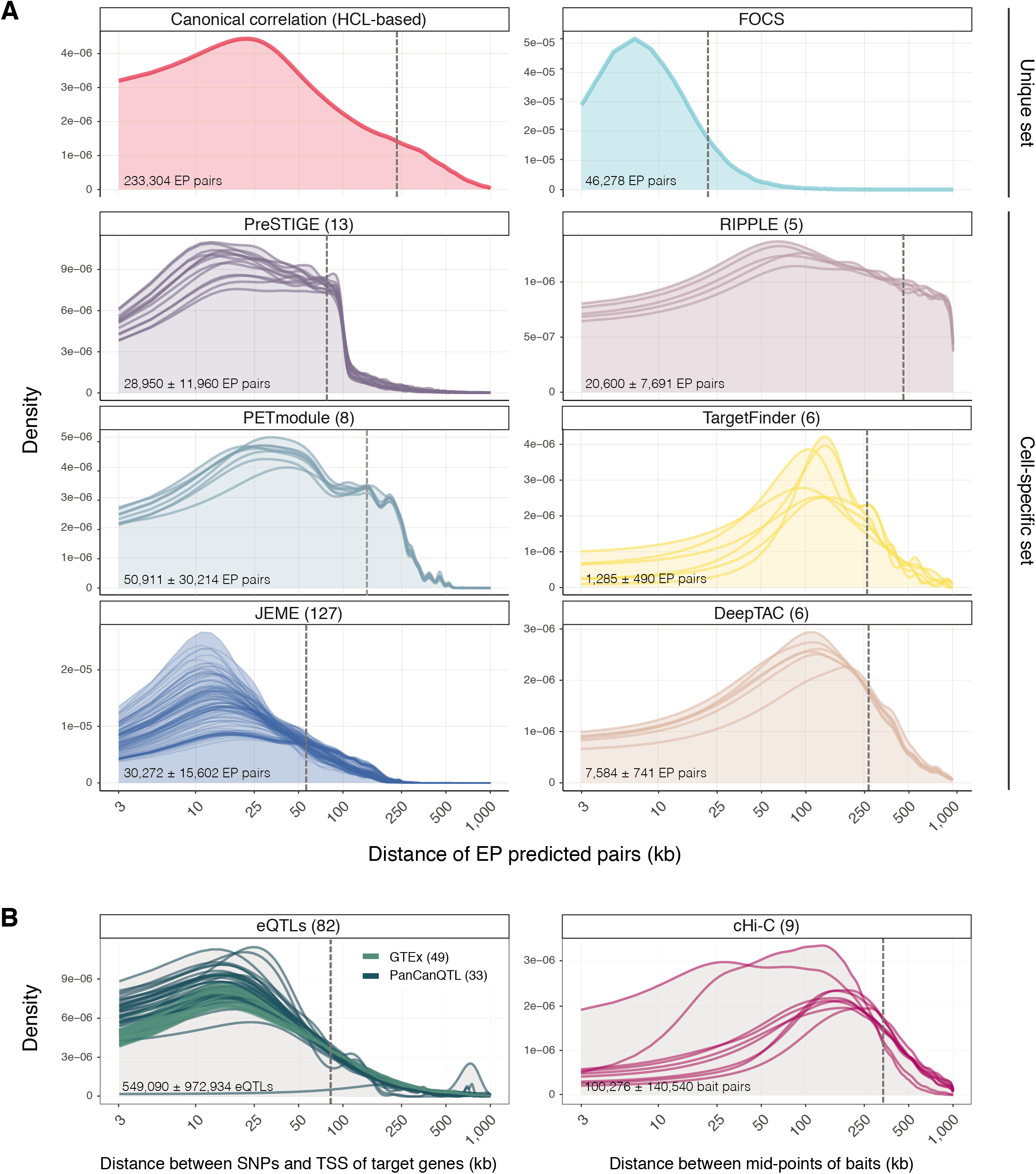
Benchmarking against other ETG pairing methods. **A**. Distributions of distances between enhancer and promoter (or TSS) of predicted pairs for our framework (HCL score > 11) and other seven ETG pairing approaches. For methods that rely on cell-type specific sets of promoters and enhancers as input, one distribution for each input set (number of input sets in brackets) is reported, along with the mean and standard deviation of the number of predicted EP pairs (text in the bottom left corner). Overall mean of distances is reported as a dotted grey vertical line. **B**. Distributions of distances of mid-range (eQTLs supported, left panel) and long-range (cHi-C supported) true positive interactions. Overall mean of distances is reported as a dotted grey vertical line. Mean and standard deviation of the number of true positive EP pairs is reported (text in the bottom left corner).

**Figure S5.**
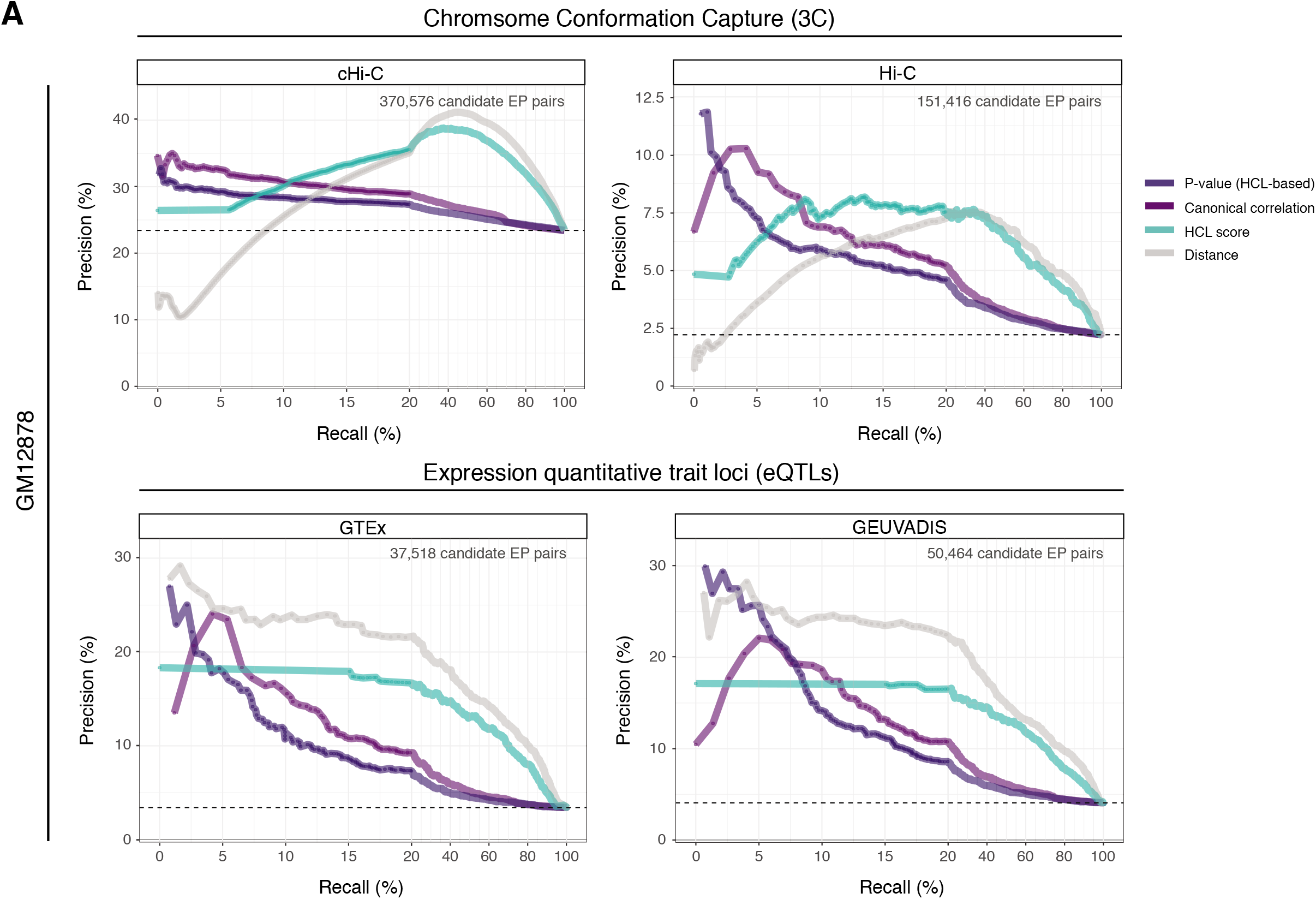
Benchmark against independent reference datasets. **A**. Precision-recall curves for GM12878 cell line BENGI benchmark interactions, assessed employing 3C supported (cHi-C, upper left panel; Hi-C, upper right panel) and eQTLs supported true (GTEx, bottom left panel; GEUVADIS, bottom right panel) ETG positive interactions. Performances are calculated for: AdaPT HCL-based adjusted p-values (dark purple lines), canonical correlation (light purple lines), HCL score (aquamarine lines) and linear EP distance (grey lines). The total number of ETG pair considered is reported on the upper right corner of each panel. The plot is reporting an expanded x-axis in the initial part of the curve (corresponding to recalls up to 20%) to provide more details in the most informative part of the chart. For higher recall values, the precision-recall curves reported here tend to converge without further crossing each other.

